# A conserved uORF regulates APOBEC3G translation and is targeted by HIV-1 Vif protein to repress the antiviral factor

**DOI:** 10.1101/2021.01.13.426487

**Authors:** Camille Libre, Tanja Seissler, Santiago Guerrero, Julien Batisse, Cédric Verriez, Benjamin Stupfler, Orian Gilmer, Romina Cabrera-Rodriguez, Melanie M. Weber, Agustin Valenzuela-Fernandez, Andrea Cimarelli, Lucie Etienne, Roland Marquet, Jean-Christophe Paillart

**Affiliations:** Université de Strasbourg, CNRS, Architecture et Réactivité de l’ARN, UPR 9002, 2 Allée Conrad Roentgen, F-67084 Strasbourg cedex, France; CIRI-International Center for Infectiology Research, INSERM U1111, Université Claude Bernard Lyon 1, CNRS, UMR5308, Ecole Normale Supérieure de Lyon, Université Lyon, F-69000 Lyon, France; Laboratorio de Inmunología Celular y Viral, Unidad de Farmacología, Sección de Medicina, Facultad de Ciencias de la Salud, Universidad de La Laguna, Campus de Ofra s/n, Tenerife, Spain

## Abstract

The HIV-1 Vif protein is essential for viral fitness and pathogenicity. Vif decreases expression of cellular restriction factors APOBEC3G (A3G), A3F, A3D and A3H, which inhibit HIV-1 replication by inducing hypermutation during reverse transcription. Vif counteracts A3G at several levels (transcription, translation and protein degradation) that together reduce the levels of A3G in cells and prevent its incorporation into viral particles. How Vif affects A3G translation remains unclear. Here, we uncovered the importance of a short conserved uORF (upstream ORF) located within two critical stem-loop structures of the 5’ untranslated region (5’UTR) of A3G mRNA for this process. A3G translation occurs through a combination of leaky-scanning and translation re-initiation and the presence of an intact uORF decreases the extent of global A3G translation under normal conditions. Interestingly, the uORF is also absolutely required for Vif-mediated translation inhibition and redirection of A3G mRNA into stress granules. Overall, we discovered that A3G translation is regulated by a small uORF conserved in the human population and that Vif uses this specific feature to repress its translation.

## INTRODUCTION

The Human Immunodeficiency virus (HIV) Vif (Viral infectivity factor) protein is essential for production of infectious particles in target cells (1). Indeed, early studies demonstrated that Vif is necessary for virus replication in primary lymphoid and myeloid cells (also called non-permissive cells) but dispensable in a subset of immortalized T cell lines (called permissive cells) (2–4). This characteristic is due to the expression of a dominant inhibitor of HIV-1 replication in non-permissive cells (5, 6), later identified as APOBEC3G (Apolipoprotein B mRNA editing enzyme, catalytic polypeptide-like 3) or A3G (7). A3G belongs to a large family of cytidine deaminases (A3A to A3H) that interfere with reverse transcription by inducing mutations during the synthesis of the viral (−) strand DNA, thus leading to cytidine to uridine transitions and production of non-infectious viral particles (8). Amongst these deaminases, A3D, A3F, A3G and A3H have been shown to efficiently block HIV-1 replication after viral entry (9–14). The HIV-1 non-structural Vif protein counteracts the antiviral activities of A3 proteins, in particular A3F and A3G, which are the most potent against HIV-1 (9, 14, 15). In the absence of Vif, A3G and A3F are packaged into viral particles (16–21) and induce viral DNA hypermutation at the next replication cycle, which in turn results in non-functional viral proteins (22). Furthermore, there is also evidence that A3G and A3F can inhibit the reverse transcription and integration steps through a deamination-independent mechanism (23–27).

In HIV-1, three distinct and mutually reinforcing mechanisms are employed by Vif to reduce A3G expression and counteract its antiviral activity (28, 29). First, Vif recruits a Cullin-RING E3 ligase 5 (CRL5) complex (composed of the Cullin 5, Elongin B/C, Rbx2, and ARIH2 proteins) to A3 proteins to promote their polyubiquitination and subsequent proteasomal degradation (30–33). This pathway was the first to be described and is well characterized (33–36). Secondly, the interaction of the transcriptional cofactor CBF-β (Core Binding Factor β) with Vif-CRL5 complex affects its association with the RUNX family of transcription factors, leading to the downregulation of RUNX-dependent genes, to which A3G belongs (37, 38). Thirdly, Vif counteracts A3G expression by reducing its translation (39–41). Indeed, we previously showed that two stem-loop structures (SL2-SL3) within the 5’-untranslated region (UTR) of A3G mRNA are essential for the translational inhibition of A3G by Vif (42). Importantly, both proteasomal degradation and translational inhibition of A3G by Vif participate to reduce the intracellular level of A3G and inhibit its packaging into viral particles, demonstrating that HIV-1 has evolved complementary mechanisms to specifically inhibit the potent antiviral activity of A3G.

Regulation of translation represents a critical layer of gene expression control, allowing rapid and localized changes in the expression of proteins in response to extra- and intracellular stimuli. Translational control can occur on a global basis by modifications of the basic translation machinery, or selectively target defined subsets of mRNAs. The latter commonly involves sequence-specific recognition of target mRNAs by trans-acting factors such as miRNA complexes or RNA-binding proteins (RBP) (43, 44). To better understand translational regulation of A3G by Vif, and the role of the two stem-loop structures within the 5’-UTR of A3G mRNA in this process, we searched for *cis*-acting regulatory elements within this region. We uncovered the importance of a short and conserved uORF (upstream Open Reading Frame) in the distal part (SL2-SL3) of the 5’-UTR of A3G mRNA. Considering that the presence of uORFs usually correlate with reduced protein expression of the downstream ORF, we studied the impact of this uORF on A3G translation and on Vif-mediated translational repression. Mutagenesis of the A3G 5’-UTR and uORF, combined with the analysis of the translation level, indicated that A3G mRNA is translated through a combination of leaky-scanning and translation re-initiation. Interestingly, disruption of the uORF abrogated the Vif-mediated downregulation of A3G translation. Furthermore, we showed that, under stress conditions, A3G mRNA was targeted into stress granules in a Vif and uORF dependent manner that correlated with A3G translation inhibition. Taken together, we discovered that human A3G translation is regulated by a small uORF embedded within its 5’-UTR and that HIV-1 Vif uses this specific feature to repress A3G translation and target it to stress granules.

## MATERIALS AND METHODS

### Plasmids

Plasmid pCMV-hA3G has been previously described (40). Mutated plasmids were generated by Quick-Change Site-directed Mutagenesis (Table 1) (Agilent Technology) based on the secondary structure model of the 5’UTR of A3G mRNA (40) and verified by DNA sequencing (Eurofins, Germany). The A3 uORF2 mutants were constructed by inserting a G after the uAUG (translation initiation codon of the uORF) and deleting a G before the uUGA (termination codon of the uORF) in order to place the uUGA in frame with the uAUG, thus changing the wild type (wt) 23 amino acid sequence MTTRPWEVTLGRAVLKPEAWSRK to MDYEALGGHFREGCPKTRSLEQK. Vif was expressed from pcDNA-hVif expression vector encoding codon-optimized NL4.3 Vif (45) or from pCRV1-B-LAI (46) expressing Vif from the pLAI.2 strain of HIV-1. Plasmids expressing stress granule (pcDNA GFP-PABP, pcDNA GFP-TIA1) or P-Body (pcDNA GFP-DCP1, pcDNA GFP-AGO2) markers were kindly provided by Dr S. Pfeffer (IBMC-CNRS, Strasbourg).

**Table 1:**
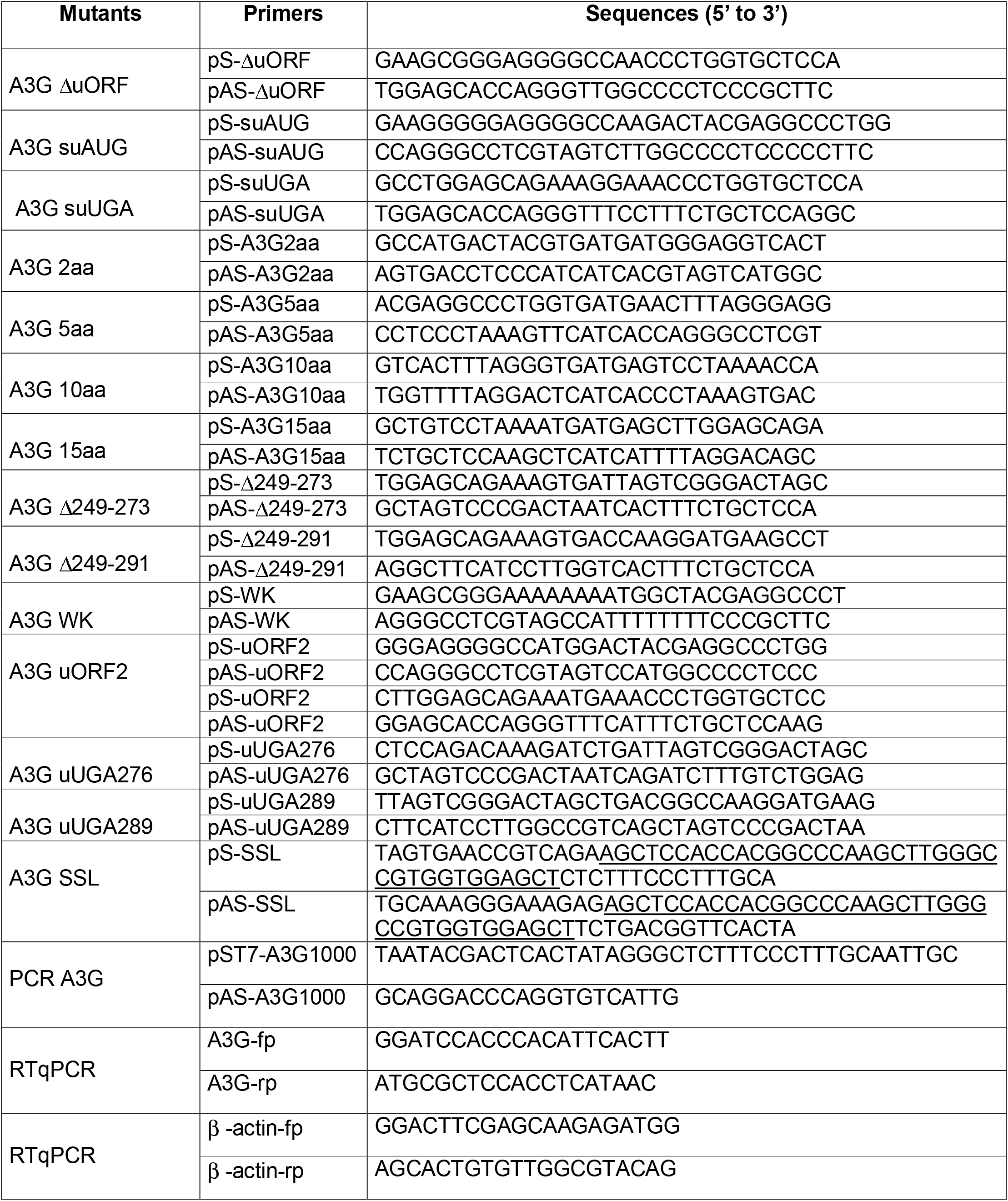
Description of the primers used in this study.

### RACE-PCR

Rapid Amplification of cDNA-ends by PCR (RACE-PCR) was performed following the instructions of the supplier in the 5’/3’ RACE Kit, 2^nd^ Generation (Roche). For the 5’-RACE-PCR, 0.2-0.5 µg of human spleen total RNA (Life Technologies) served as template to synthesize the cDNA corresponding to the 5’-end of A3 RNAs by using the Transcriptor Reverse Transcriptase and a specific primer 1 (SP1) according to manufacturer recommendations. The cDNAs were produced for 1 h at 55°C and the reaction was stopped by heating the mixture at 85°C for 5 min. After a purification step, a poly-A tail was added to cDNAs, which were then amplified by a second PCR. This PCR used a dT-Anchor Primer and a second SP2 (0.25 µM each) to amplify 5 µl of polyadenylated cDNAs in a 50 µl mix containing 1 U of Phusion Polymerase, 0.2 mM dNTPs and 1.5 mM MgCl_2_. The PCR protocol was the following: 3 min at 98°C and 10 cycles of 15 s at 98°C, 30 s at the optimal annealing temperature and 1 min at 72°C. The elongation step was then increased by 20 s steps until it reached 2 min and 23 cycles were performed. A final elongation step of 7 min ended the amplification. A nested PCR was performed with 1 µl of amplicons from the last PCR, in an identical reactional mixture, except for the SP3 used. PCR products were cloned into pJET 1.2 vector (Thermofisher) and sequenced.

For the 3’-RACE-PCR, as mRNAs for the total RNA extract were already polyadenylated, the purification and poly-A tailing steps were not necessary. Using 1 µg of total RNA and a dT-Anchor primer, the cDNAs were synthesized according to the manufacturer protocol and used for a PCR amplification with a PCR Anchor Primer and SP5 primer. Both PCRs were performed as described above. A nested PCR, absent from the initial protocol, was added with a SP6 primer and the same PCR Anchor Primer used in the last PCR to obtain enough material for bacterial transformation. PCR products were cloned into pJET 1.2 vector (Thermofisher) and sequenced.

### RNA modification by SHAPE (selective 2’-hydroxyl acylation analyzed by primer extension)

#### RNA synthesis

RNAs corresponding to the first 1,000 nucleotides of A3G mRNAs were synthesized by *in vitro* transcription from PCR products obtained from wt and mutant pCMV-hA3G plasmids ((40) and this study). PCR was performed using primers pST7-A3G1000 and pAS-A3G1000 (Table 1) with the following protocol: 3 min at 95°C and 34 cycles of 30 s at 95°C, 30 s at 56°C and 1 min at 72°C, with a final elongation step of 5 min at 72°C. Three additional Gs were inserted immediately downstream the T7 promotor (5’-end of RNA) to increase transcription efficiency. Transcription was performed using a MEGAscript T7 Transcription kit (Thermo Fisher Scientific) following the manufacturer’s instructions and purified by exclusion chromatography on a TSKgel G4000SW column as previously described (40). RNA integrity and purity were confirmed by denaturing polyacrylamide gel electrophoresis.

#### RNA modification with NMIA

2 pmol of A3G RNA was heated at 90°C for 2 min and placed on ice for 2 min, then refolded 20 min at 37°C in 20 µl of HEPES buffer (final concentration: 30 mM HEPES pH 8.0 ; 300 mM KCl ; 5 mM MgCl_2_), then 1 µg of a tRNA mix was added (total volume adjusted to 30 µl), and refolding further continued 10 min at room temperature. One pmol of RNA was then treated in a final volume of 18 µl with 10 mM NMIA (Sigma-Aldrich) in anhydrous DMSO (Sigma-Aldrich) (+) or only with anhydrous DMSO (control (−)). After 50 min at room temperature, deionized water was added and material was precipitated with 3 volumes of 100% ethanol, 1/10 volume of 3 M sodium acetate pH 5.0, 1 µl of glycoBlue (ThermoFisher) for 30 min in a dry ice/ethanol bath and collected by centrifugation at 20,800 g for 30 min at 4°C. The pellets were washed twice with 70% ethanol, dried and resuspended in deionized water (7 µl final volume).

#### cDNA synthesis and analysis by capillary electrophoresis

RNA samples treated with NMIA (+) or DMSO (−) were mixed with 1 pmol of AS primer 1 (5’-TGGGTGGTACTTAAGTTCGG-3’: complementary to nts 476-495 of A3G RNA) labeled with Vic (Life Technologies SAS, France). The mixture was heated at 90°C for 2 min and placed on ice for 2 min. After the addition of 2 µl of avian myeloblastosis virus (AMV) reverse transcriptase (RT) Buffer (final concentration: 50 mM Tris-HCl pH 8.3 ; 100 mM KCl ; 4 mM DTT ; 10 mM MgCl_2_) and 10 min incubation at room temperature, reverse transcription was performed by adding 2 µl of AMV RT buffer (Life Sciences), 6 µl dNTPs 2.5 mM (Invitrogen), and 2 U of AMV RT (Life Sciences) in a final volume of 20 µl. Elongation was ensured by incubation 20 min at 42°C followed by 30 min at 50°C. The enzyme was inactivated at 60°C for 10 min. Simultaneously, a sequencing reaction was performed with 2 pmol of unmodified RNA and 2 µl of a 2 mM AS primer 1 labeled with Ned (Life Technologies SAS, France). Reverse transcription was performed as for the SHAPE (+) and (−) elongation reactions except for the nucleotide mix added, which was composed of 6 µl G10 (0.25 mM dGTP, 1 mM dATP, 1 mM dCTP, 1 mM TTP), 2 µl ddGTP at 100 µM, 2 µl AMV RT buffer and 2 U of AMV RT (Life Sciences).

The reaction volumes were adjusted to 100 µl and cDNAs were phenol-chloroform extracted (Roti-phenol). For each experiment, the modified (+) and unmodified (−) samples were pooled each with a ddG sequencing reaction before ethanol precipitation. The cDNAs were resuspended in 10 µl HiDi Formamide (Applied Biosystem), denatured at 90°C for 5 min, placed on ice for 10 min and finally centrifuged for 5 min at 6,000 g. The primer extension products were loaded on an ABI 3130XL Genetic Analyzer (Applied Biosystem) and the electropherograms were analyzed with the QuShape software to extract reactivity data for each sample. The mean reactivity data from at least three independent experiments were obtained for each sample and used as constraints to fold the RNA secondary structure with the RNAstructure software version 6.0 (47). No other constraints than the SHAPE reactivities were applied to the fold. The dot bracket file obtained from RNAstructure was then used to translate the structural data into VARNA version 3–93 (48) where the validated structure was redrawn.

### Cell culture

HEK 293T cells were maintained in Dulbecco’s modified Eagle’s medium (DMEM, Life Technologies) supplemented with 10% fetal bovine serum (PAA) and 100 U/ml penicillin/streptomycin (Life Technologies) at 37°C and 5% CO_2_ atmosphere. Transfections of HEK 293T cells were carried out using the X-tremeGene 9 DNA Transfection Reagent (Sigma-Aldrich) as recommended by the manufacturer. Briefly, 700,000 cells/well were seeded at 70% confluence in a 6-well plate and co-transfected with 100 ng of pCMV-hA3G constructs and 1 µg of pcDNA-hVif. Cells were also exposed to the chemical proteasome inhibitor ALLN (25 µM) for 14 h.

For FISH and immunofluorescence (IF) experiments, cells were plated on a glass coverslip in 6 well-microplates (700,000 cells/well for HEK 293T cells and 350,000 cells/well for HeLa cells). Cells were cultured in DMEM. Transfections of HEK 293T and Hela cells were carried out as described above. When required, 0.5 µg of stress granules (GFP-PABP or GFP-TIA1) or P-Bodies (GFP-DCP1 or GFP-AGO2) markers were transfected. Twenty-four hours post transfection, stress induction was performed either by treating cells with 500 μM sodium arsenite (NaAsO_2_, Sigma-Aldrich) or by incubating them at 44°C (heat shock) for 30 min before further analysis.

### Immunoblotting

Twenty-four hours post-transfection, cells were washed in 1X PBS (140 mM NaCl, 8 mM NaH_2_PO_4_, 2 mM Na_2_HPO_4_) and lysed for 10 min at 4°C in 1X RIPA (1X PBS, 1% NP40, 0.5% sodium deoxycholate, 0.05% SDS) supplemented with protease inhibitors (complete EDTA Free cocktail, Roche). After 1 h centrifugation at 20,817 g, cell lysates were adjusted to equivalent protein concentration (Bradford assay, Bio-Rad), fractionated on Criterion TGX 4-15% gels (Bio-Rad) and transferred onto 22 µm PVDF membranes using the Trans-Blot Turbo™ Transfer System (Bio-Rad). Blots were probed with appropriate primary antibodies. Polyclonal anti-A3G (#9968), anti-A3H (#12155), anti-A3F (#11226) and monoclonal anti-Vif (#319) antibodies were obtained through the NIH AIDS Research and Reference Reagent Program. Polyclonal anti-A3C (#EB08307) and anti-A3D (#GTX87757) antibodies were obtained from Everest Biotech and Genetex, respectively. Monoclonal anti-β-actin antibody was purchased from Sigma-Aldrich (#A5316). The PVDF membranes were incubated with horseradish peroxidase-conjugated secondary antibodies (Bio-Rad), and the proteins were visualized by enhanced chemiluminescence (ECL) using the ECL Prime Western blotting detection reagent (GE Healthcare) and the ChemiDoc™ Touch Imaging System (Bio-Rad). Bands were quantified using Image J. Student’s T-test was used to determine statistical significance.

### Real-time qPCR

Twenty-four hours post-transfection, total RNA was isolated from HEK 293T cells using RNAzol®RT (Euromedex). After RNase-free DNase treatment (TURBO DNA-free kit, Invitrogen), total RNA (1 µg) was reverse-transcribed using the iScript™ Reverse Transcription Supermix (Bio-Rad) as recommended by the manufacturer. Subsequent qPCR analysis was performed using the Maxima™ SYBR Green qPCR Master Mix (ThermoFisher) and was monitored on a CFX Real Time System (Bio-Rad). Gene-specific primers for A3G and β -actin are detailed in Table 1. The A3G mRNA levels were normalized to those of actin mRNA and relative quantification was determined using the standard curve-based method.

### FISH and immunofluorescence (IF) assays

The FISH (Fluorescence In Situ Hybridization) probe was obtained as follow: fragment from nucleotide position 100 to 406 of A3G mRNA was cloned between EcoRI and Xba1 sites into a pcDNA vector. After linearization by XbaI, T7 in vitro transcription was performed in presence of conjugated DIG-11-UTP (Roche) following the manufacturer instruction (1 mM of each dNTP, except dUTP at 0.65 mM and 0.35 mM of the labeled DIG-11-UTP). After DNAse I (Roche) treatment, A3G specific probe was purified by phenol-chloroform extraction and ethanol precipitation. Pellets were resuspended in 50 µl of deionized water giving a 100X concentrated A3G mRNA probe. Aliquots of 1 µl were stored at − 80°C.

For FISH and IF (Immuno-Fluorescence) assays, cells were fixed with 4% (w/vol) paraformaldehyde/PBS for 20 min at RT. Fixation was stopped in 100 mM glycine for 10 min at room temperature. Cells were then permeabilized with 0.2% (w/vol) TRITON X-100/PBS solution at room temperature for 5 min. For FISH assays, after 2 washes in PBS, coverslips were treated for 15 min at room temperature with DNAse I (Roche – 25 U/coverslip) then washed with 1 ml of PBS. One µl of specific A3G mRNA probe was diluted 100x in deionized water (concentration ∼ 5 ng/µl). Coverslips were loaded with 50 µl of pre-warmed hybridization solution (formamide 50%; tRNA 0.1 µg/µl; SSPE 2X (NaCl 300 mM, NaH_2_PO_4_ 20 mM, EDTA 2 mM); Denharts solution 5X (Ficoll 0.1%, Polyvinylpyrrolidone 0.1%, BSA 0.1%); RNAse OUT 0.1 U/µl and 25 ng of specific probe) for 16 h at 42°C, in humid atmosphere. Coverslips were then washed with 50 µl of pre-warmed buffer containing 50% formamide and SSPE 2X for 15 min at 42°C, and then twice with 50 µl SSPE 2X for 5 min at 42°C. Finally, for both FISH and IF assays, after a wash in PBS, coverslips were blocked with 3% (w/vol) BSA in PBS for 1 h at room temperature. Primary antibodies were diluted in 3% (w/vol) BSA/PBS and incubated 3 h at 37°C, followed by incubation of secondary antibodies at room temperature for 2 h in the dark. After a brief wash in PBS, coverslips were mounted in one drop of SlowFade gold antifade reagent (Thermo Fisher) with (IF) or without (FISH) DAPI (Thermo Fisher). The following primary antibodies were used: sheep polyclonal anti-DIG (Roche); Rabbit polyclonal anti-hA3G-C17 (#10082) and mouse monoclonal anti-HIV-1 Vif (#319) antibodies were obtained through the NIH AIDS Research and Reference Reagent Program. All were used at a 1:500 dilution. The following dye-conjugated secondary antibodies were used: Alexa Fluor 647 anti-rabbit, Alexa Fluor 647 anti-goat, Alexa Fluor 488 anti-mouse, Alexa Fluor 546 anti-sheep, Alexa Fluor 405 anti-mouse, Alexa Fluor 594 anti-goat (Life Technologies). All were used at a 1:200 dilution.

### Microscopy and image analysis

Images were acquired on a Zeiss LSM780 spectral microscope running Zen software, with a 63x,1.4NA Plan-Apochromat oil objective at the IBMP microscopy and cellular imaging platform (Strasbourg, France). Excitation and emission settings were spectrally selected among the 4 laser wavelengths available in the microscope (405; 488; 561; and 633 nm) according to the secondary antibody used. Image processing (contrast, brightness and merges) were performed with ImageJ 1.43m software (49). An additional macro allowing the multichannel profile plot has been designed by Jerome Mutterer from the Microscopy platform. Percentage of cells showing a co-localization between FISH signal (A3G mRNA) and stress or P-Body markers were counted on subsets of 100 cells (n=3).

### APOBEC3 genotype data

The NCBI dbSNP database (https://www.ncbi.nlm.nih.gov/snp, July 31^st^ 2019) as well as the UCSC Genome Browser database (https://genome.ucsc.edu/, July 31^st^ 2019) were mined for polymorphic variants in human A3G and A3F mRNA 5’UTRs.

## RESULTS

### In silico analyses of A3G mRNA 5’UTR identify a conserved uORF

To understand the mechanism by which Vif inhibits A3G mRNA translation in a 5’-UTR-dependent manner (40, 42), we searched for *cis*-acting elements within the 5’-UTR that could contribute to this repression. Using a computational platform capable of identifying RNA regulatory elements (RegRNA 2.0) (50), we identified a terminal oligopyrimidine (TOP) element that we already excluded from Vif-mediated translational control of A3G mRNA as the deletion of the 5’-end region (mutant A3G SL2-SL3) did not impact the Vif-mediated translation inhibition (42), and an uORF within the 5’-UTR of A3G mRNA sequence (Figure 1). uORFs are cis-regulatory elements that negatively regulate translation of downstream ORF (51–54). In A3G mRNA, our analysis showed that an uORF is encoded within the SL2 and SL3 domains of the 5’-UTR (Figure 1). Of note, we previously showed that these two SLs are required for Vif-mediated A3G translation inhibition (42), suggesting this uORF could be involved in translational inhibition. The uORF is positioned between nucleotides 177 (initiation codon uAUG) and 248 (termination codon uUGA) within the 5’-UTR, 49 nucleotides upstream of the main AUG (mAUG) initiation codon of A3G (nucleotide 298). It encodes a putative peptide of 23 amino acids (Figure 1) without any particular motif according to the PROSITE database (https://prosite.expasy.org). We also observed that the Kozak contexts around the upstream and major AUGs are considered as adequate for translation initiation, suggesting the uAUG is functional (55). The Kozak consensus is NNN*R*NN<u>AUG</u>(*A/C/U*) where N is any base, *-3R* is a purine, *+4* position is A, C or U (the translation codon is underlined) and the actual sequences are GGG*G*CC<u>AUG</u>*A* and GCCA*A*GG<u>AUG</u>*A* for the uAUG and mAUG, respectively (the important −3 and +4 positions are in italic). The same uORF is also present in A3F mRNA, with 94% and 96% identity at the nucleotide and amino acid level, respectively. To determine if these uORFs are conserved in the human population, we mined the NCBI dbSNP (single nucleotide polymorphism database). In the A3G uORF (+/− 10 nucleotides) region, which included the Kozak context, we did not find any variant with a Minor Allele Frequency (MAF) above 0.005 (Table S1). In the A3F uORF region, there was a single common SNP (MAF>0.01, rs35898507) that would induce an amino acid change from G to V in the uORF (G being the ancestral allele present at >96% in the population) (Table S1). Overall, we did not find any variant with a MAF>0.0005 that would dramatically impact the uORFs of A3G and A3F (i.e. that would impact the start codon, the stop codon, or would induce a frameshift). Lastly, the genetic distance between the uORF and the mAUG was also conserved. This shows that the herein identified A3G and A3F uORFs are highly conserved in the human population.

**Figure 1:**
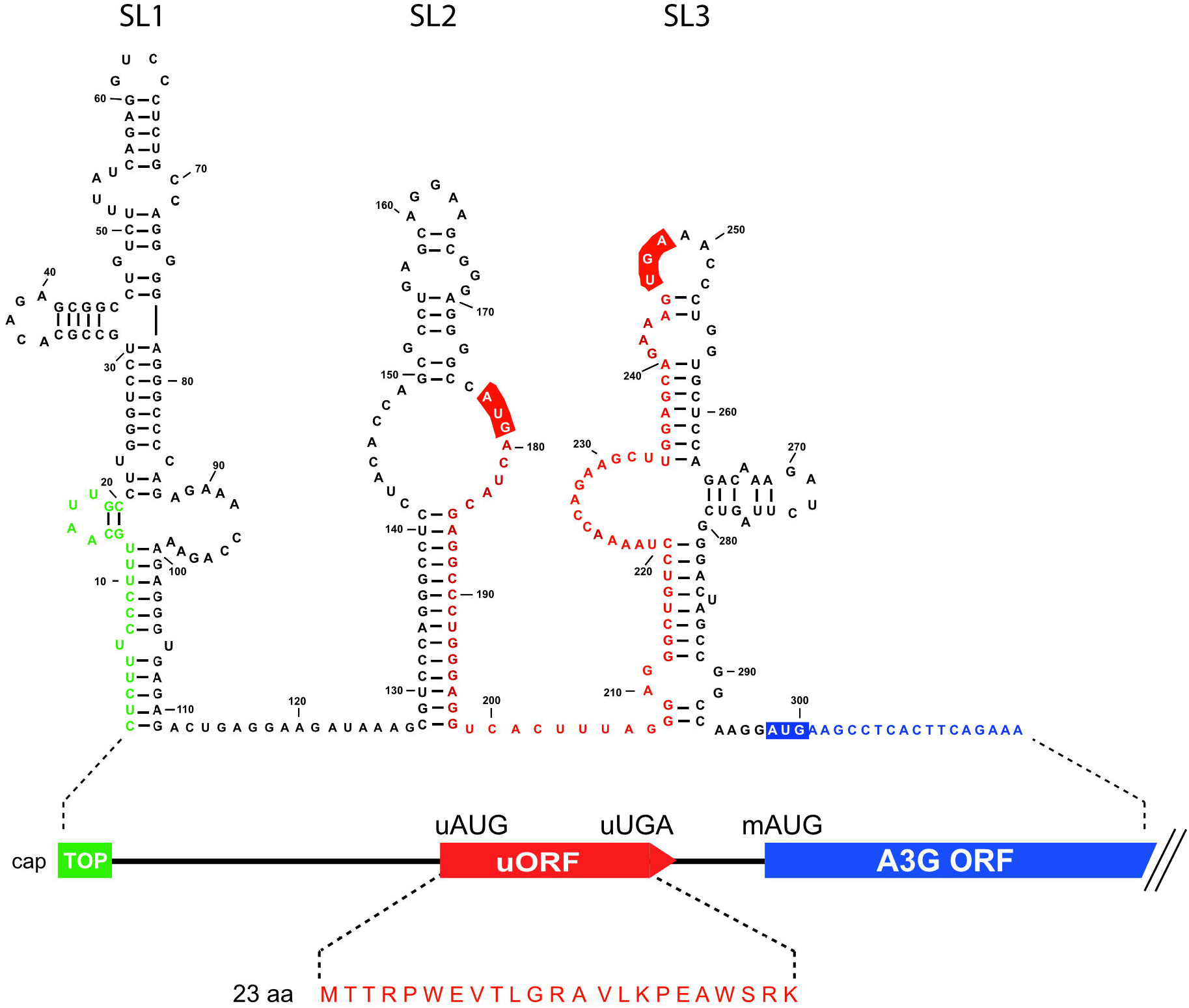
Schematic representation of the 5’-UTR of A3G mRNA. The secondary structure of the 5’-UTR of A3G mRNA as determined in (40) is indicated, as well as the TOP element (green) and of the uORF (red). The putative peptide expressed from the uORF is also indicated.

### Vif specifically regulates the translation of A3G and A3F among members of the A3 family

We previously showed that A3G was regulated at the translational level by HIV-1 Vif (40, 42). We now wonder whether this translational regulation could be extended to other A3 proteins. Thus, we performed 5’-and 3’-RACE (5’-rapid amplification of cDNA ends analysis) from human spleen total RNA with A3B, A3C, A3D and A3H specific primers (Table 2) in order to identify their 5’- and 3’-UTRs (A3A vector (NM_145699.3) was a generous gift from Dr Vincent Caval, Pasteur Institute, Paris (56); A3F (NM_145298.5) and A3G (NM_021822) were already available in our lab). After cloning and sequencing of the expression constructs, corresponding full-length mRNA sequences were synthesized (Proteogenix, France) and consequently cloned into pCMV vectors (Figure 2A). The 5’-UTR sequences comprised 44, 106, 407 and 127 nucleotides for A3B, A3C, A3D and A3H, respectively, while their 3’-UTR sequences contained 572, 431, 934 and 372 nucleotides, respectively (Figure 2A and Table S2). Sequences obtained for the 5’-UTR of A3D and A3H correspond to those published in GenBank™ (accession number NM_152426.3 and NM_001166003.1, respectively). The 5’-UTR sequence of A3B is shorter (44 versus 55 nucleotides) than the reference (NM_004900.4), while the one from A3C is a bit longer (106 versus 103 nucleotides) than the reference (NM_014508.2). All A3 clones were transfected into HEK 293T cells in presence or absence of Vif and with or without ALLN (proteasome inhibitor) to discriminate between translational inhibition and proteasomal degradation (42). Our results showed that all A3 proteins are well expressed from full-length mRNA expression constructs (Figure 2B, lanes 2) and are degraded in response to Vif expression as expected (Figure 2B, lanes 3 and Figure 2C, red bars), except for A3A and A3B. This behavior is not unusual for A3A/A3B and could be linked to their Vif-induced nucleocytoplasmic transport (57). A3A presents two isoforms (A3A-L and A3A-S) due to a leaky scanning mechanism as previously observed (56). A3D was not significantly reduced as previously observed (58). However, in presence of ALLN, we only observed a significant decrease in A3G and A3F expression when Vif was present (Figure 2B, compare lanes 4 & 5; Figure 2C blue bars), suggesting that these two A3 proteins are the only ones to be regulated at the translational level by Vif. Moreover, sequence analysis of the 5’-UTR of A3A, A3B, A3C, and A3H did not reveal the presence of any uORF or other regulatory motifs. A short putative uORF (30 nucleotides) is present within the 5’-UTR of A3D (Table S2) but was not involved in the Vif-mediated translational inhibition (Figure 2B and 3C).

**Table 2:**
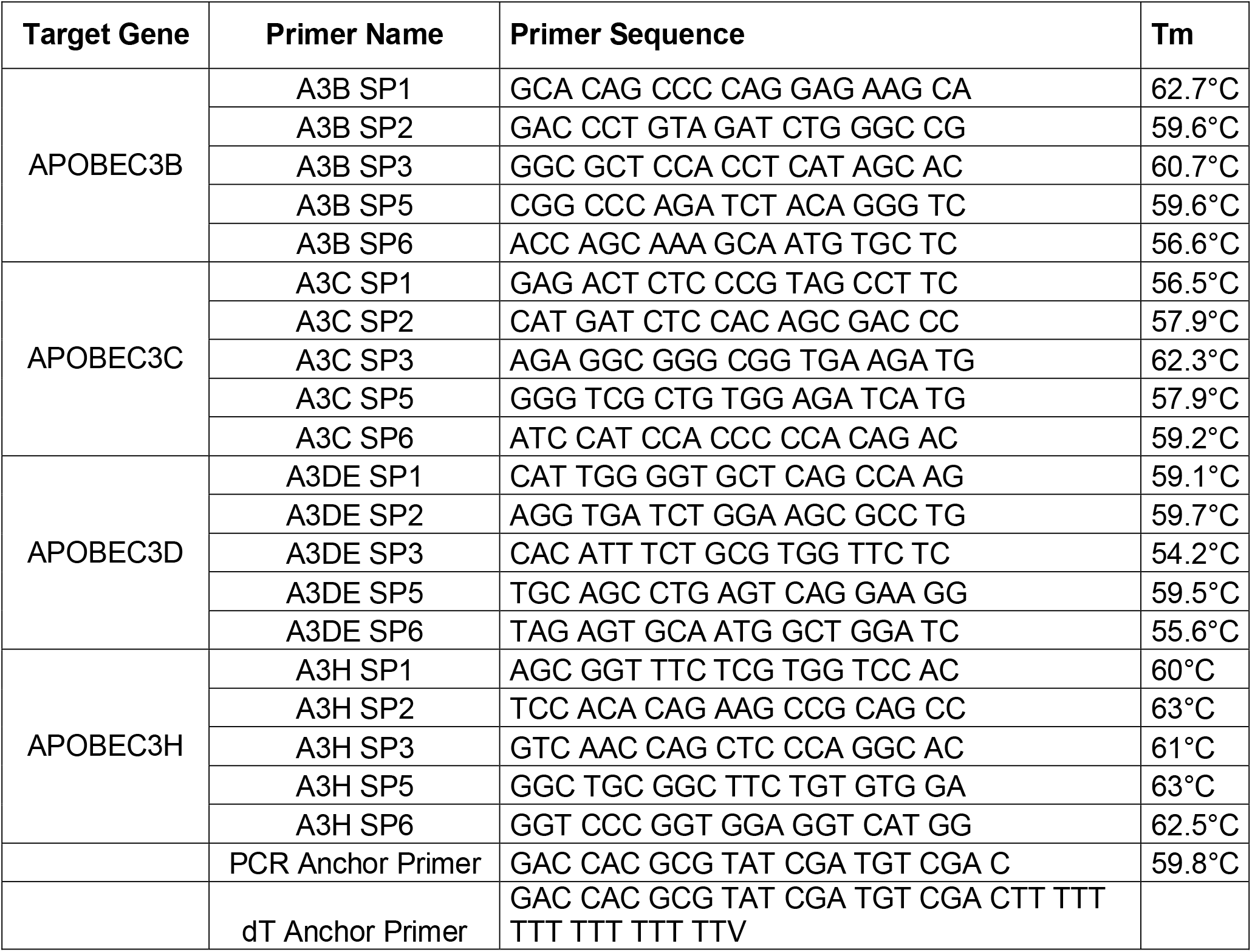
Description of the primers used for the RACE PCRs.

**Figure 2:**
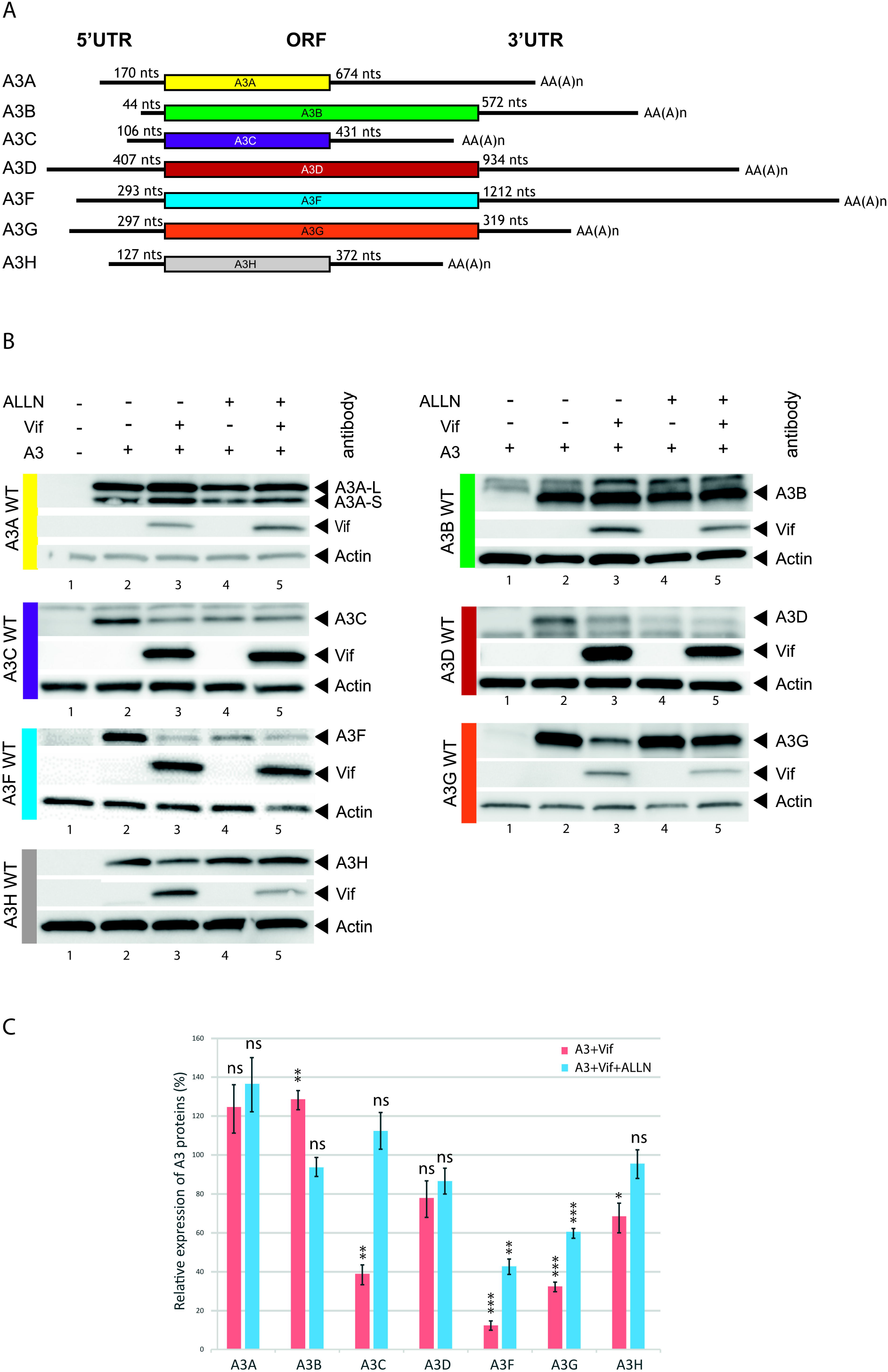
Vif does not inhibit the translation of all APOBEC3 mRNAs. **A**) Schematic representation of A3 mRNAs identified by RACE-PCR. The lengths of the various 5’- and 3’-UTRs are indicated. **B**) HEK 293T cells were transfected with plasmids expressing the different A3 proteins in the presence or absence of Vif and in the presence or absence of a proteasome inhibitor (ALLN). Proteins were separated by SDS/PAGE and analyzed by immunoblotting with appropriate antibodies (see Material and methods) except A3A and A3B which were detected with anti-A3G (#9968) antibody. **C**) Bands were quantified using Image J and relative expression of A3 proteins is represented. After normalization to actin, A3+Vif (red bars, lanes 3 of panel B) and A3G+Vif+ALLN (blue bars, lanes 5 of panel B) were compared to their equivalent without Vif (lanes 2 and 4, respectively) set to 100%, respectively. Data represent the mean ± S.E.M. for at least three independent experiments. P-values are indicated as follows: *<0.05; **<0.01; ***<0.001. ns (not significant): > 0.05.

### Mechanism of A3G mRNA translation

Since uORFs usually reduce translational efficiency by approximatively 30-50% (54), we asked whether this motif naturally participates in A3G mRNA expression. First, we inactivated the uORF of this mRNA by either deleting the entire uORF (A3G ΔuORF), or by substituting the uAUG (A3G suAUG) (Figure 3A). The expression of these mutants was examined after transfection of HEK 293T cells and immunoblotting against A3G protein (Figure 3B). As expected, uORF inactivation significantly increased A3G protein expression (Figure 3B, compare lanes 2-3 to lane 1), suggesting the uORF intrinsically represses A3G mRNA translation. Moreover, we tested whether those mutations affect mRNA levels by RTqPCR and showed that A3G suAUG mRNA level was reduced by 30% (Figure 3B, lane 3 blue bar), further supporting an increase of A3G translation (ratio protein/RNA=1.8 and 1.4 for A3G suAUG and A3G ΔuORF, respectively).

**Figure 3:**
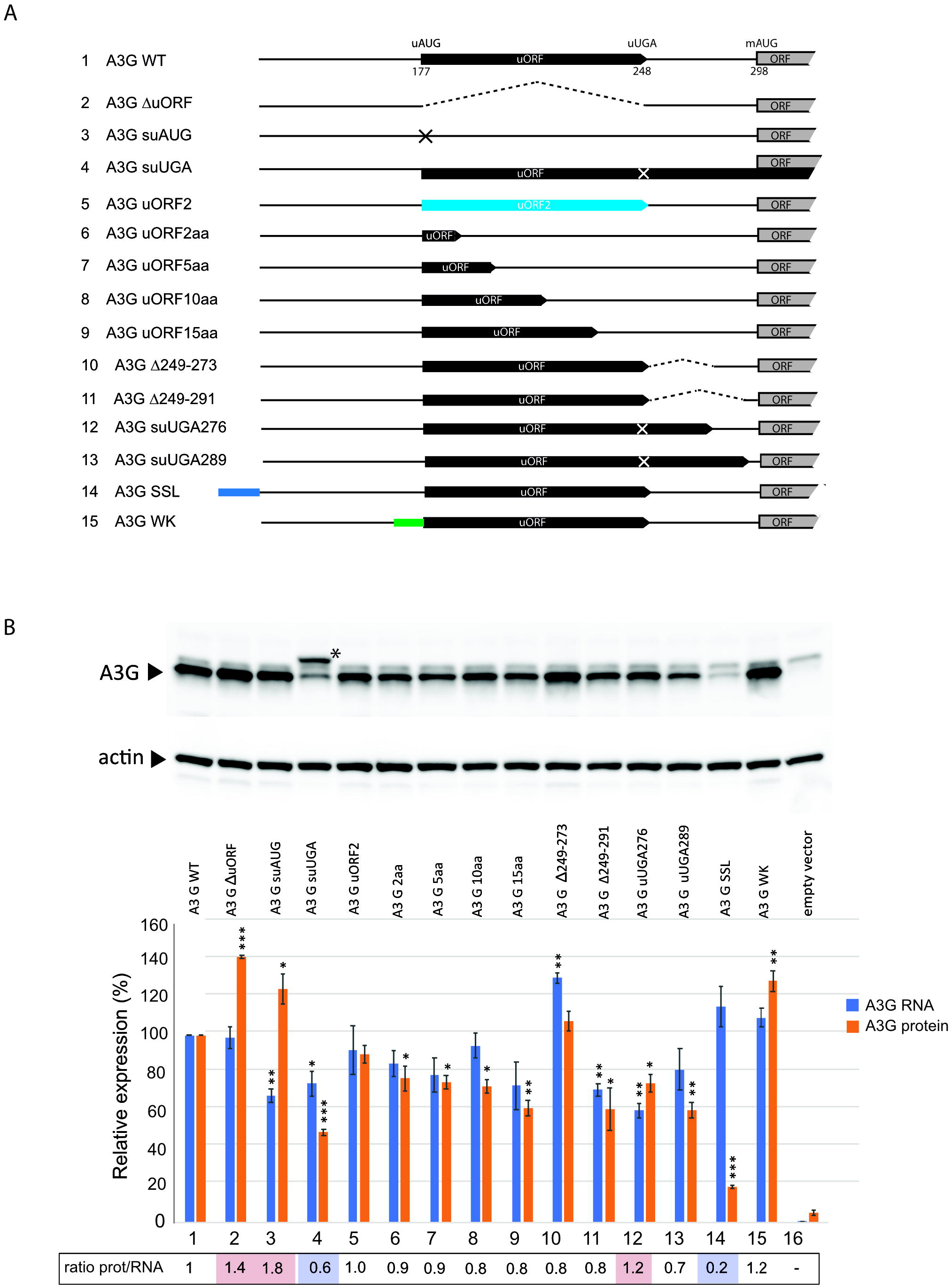
Importance of the uORF for A3G mRNA expression. **A**) Schematic representation of the different A3G 5’-UTR constructs used in this study. **B**) HEK 293T cells were transfected with wt or mutated A3G constructs. Proteins were separated by SDS/PAGE and analyzed by immunoblotting with anti-A3G (#9968) and anti-β-actin (#A5316) antibodies. Bands were quantified using Image J, normalized to actin, and relative expression of A3G is represented in a histogram (orange bars). Total RNA was extracted from transfected cells and RTqPCR were performed to study the relative expression of wt and mutated A3G constructs (blue bars). The ratio protein/RNA is indicated below. Data represent the mean ± S.E.M. for at least three independent experiments. P-values are indicated as follows: *<0.05; **<0.01; ***<0.001.

The results presented above show that the uORF negatively regulates A3G translation. According to the literature (51–54, 59), uORFs can regulate the translation of the main ORF by different mechanisms, in cis or in trans through the encoded peptide, or by affecting leaky-scanning, direct translation initiation at an IRES or translation re-initiation. To analyze the mechanism of A3G translation, we constructed a series of A3G mRNAs with mutations in the uORF sequence or its surrounding nucleotides and analyzed their expression after transfection of HEK 293T cells (Figure 3).

First, we tested the possibility that the uORF peptide acts in *trans* to regulate the main ORF translation. To achieve this, we changed the uORF amino acid sequence by shifting the uORF by one nucleotide (see Materials and methods) and studied its effect on A3G expression (mutant A3G uORF2). We observed no effect of the putative peptide in A3G expression (Figure 3B, lane 5), consistent with the fact that peptides expressed from an uORF generally do not play a functional role in translation regulation (60, 61). This result also suggests that the function of the uORF is dependent on features that drive uORF translation rather than the specific peptide produced.

Next, we examined if translation initiation at the uAUG is required for the regulatory mechanism. The frequency of ribosomal recognition of a translation initiation codon is determined by its sequence context (62), and positioning of a translation initiation codon within a “poor” sequence context will result in inefficient ribosomal recognition and bypassing (leaky scanning). We thus replaced the uAUG Kozak consensus sequence (GGGGCC<u>AUG</u>A) by a weak non favorable context (GAAAAA<u>AUG</u>A, mutant A3G WK) (Figure 3A). Consistent with the predicted decrease in recognition of the modified uAUG context by ribosomes, we observed a significant increase in protein expression (Figure 3B, lane 15), suggesting that the natural uAUG Kozak context is an essential element for the translational repression.

In a natural context of A3G mRNA, the uORF encodes a putative peptide of 23 amino acids. Next, we created a mutant RNA where the translation termination codon of the uORF (uUGA) is inactivated and the uAUG placed in frame with the downstream major ORF (mAUG), thereby producing a fusion protein formed by the peptide produced from the uORF, the intercistronic region and the A3G coding sequence. Thus, this construct (A3G suUGA) allows to directly monitor upstream translation initiation, prevents uORF termination and, hence re-initiation, and downstream translation is only possible by leaky scanning or through a direct entry of the ribosome (IRES - Internal Ribosome Entry Site) at the mAUG. Interestingly, two protein bands can now be observed (Figure 3B, lane 4). Indeed, in addition to the standard A3G protein encoded by the main ORF (Figure 3B, lane 4, lower band - 384 amino acids), a N-terminally elongated version (424 amino acids) produced by translation initiation at the uAUG can be detected (Figure 3B, lane 4, asterisk), demonstrating that the pre-initiation complex assembled on the uAUG. Moreover, we observed a significant decrease in the expression of the main A3G protein for this mutant (about 50%, with a ratio prot/ARN: 0.6) (Figure 3B, lane 4). Since re-initiation is impossible in this case due to the mutated uORF stop-codon, these results strongly suggest that, in the wt situation, A3G is partially expressed by ribosomes re-initiating at the mAUG after having translated the uORF. However, the fact that wt A3G protein is still detected in this mutant also indicates that translation of the main ORF also involves leaky-scanning or IRES-dependent translation.

To test the IRES hypothesis, we inserted a stable stem-loop at the 5’-end of A3G mRNA (Figure 3A, mutant A3G-SSL) that prevents binding of the 40S ribosomal subunit and inhibits ribosome scanning (63). Then, if an IRES is present within the 5’-UTR of A3G mRNA, ribosomes should load directly on this site and efficiently initiate translation. As observed in Figure 3B (lane 14), expression of this mutant was very low (20%, ratio prot/ARN: 0.2), ruling out the hypothesis of an efficient IRES-dependent A3G translation.

To further analyze re-initiation, we constructed additional mutants based on previous reports (64, 65). Indeed, re-initiation is more efficient when uORF sequences are short (65) whereas leaky-scanning is not dependent on the length of the uORF (64). Then, reducing the uORF length should enhance A3G expression if a re-initiation mechanism is involved. We thus tested mutants of A3G uORF where the putative 23 amino acids peptide was reduced to 15, 10, 5 and 2 amino acids (Figure 3A). The results showed that these mutants significantly decreased (20-40%, ratio prot/RNA: 0.8-0.9) A3G protein expression (Figure 3B, lanes 6-9), suggesting that reducing the length of the uORF somehow down-regulates the translation of A3G mRNA. Because re-initiation is also dependent on the distance between the stop-codon uUGA and the main AUG of an ORF (inter-ORF region) (64), we then reduced this distance to 24 (mutant A3G Δ249-273) and 6 (mutant A3G Δ249-291) nucleotides by deletions, or by inserting a stop codon at positions 276 (A3G suUGA276) or 289 (A3G suUGA289), thus reducing the inter-ORF region to 21 and 9 nucleotides, respectively (Figure 3A). If A3G translation initiation is mediated by a re-initiation mechanism, reducing this distance should lower protein expression. Interestingly, A3G protein expression was significantly decreased (ratio prot/RNA: 0.7-0.8) when the inter-ORF region was reduced (figure 3B, lanes 10-13), suggesting that A3G may also be translated through a re-initiation mechanism, except for mutant A3G suUGA276 (ratio prot/RNA: 1.2), which did not show any significant effect on A3G protein expression (figure 3B, lane 12).

To exclude potential structural effects of the mutations on A3G translation, we analyzed the secondary structure of wt and mutant A3G mRNAs by SHAPE (see Material and methods). This analysis was performed on *in vitro* transcribed RNA corresponding to the first 1,000 nucleotides of A3G mRNA. Here, we focused on the 5’-UTR (nucleotides 1-300) as it contains all mutations described in this study. First of all, we confirmed that the 5’-UTR of wt A3G mRNA folds into independent domains that are mostly similar to our previously published secondary structure (40), with three main stem-loop structures (SL1, SL2 and SL3) and the uORF encompassing SL2 and SL3 (Figure S5 and Table S3 for the reactivity values). Differences between wt structures are mainly due to the size of RNA that was used in both studies (1,000 nucleotides versus 300 in (40)). Of note, domain 4 (SL4), which contains the mAUG, was not previously predicted as it was not included in our RNA (40), and these four SLs are connected through a stem involving long-distance interactions between regions 98-122 and 381-405 (Figure S5). Interestingly, most of tested A3G mRNA mutants contain the four mentioned domains (Figures S5 and S6). The only exceptions are the two mutants containing a large deletion (A3G ΔuORF and A3G Δ249-289), which despite a conservation of SL1, SL2, and the interdomain stem, showed structural perturbation of domain 3 and/or 4 (figure S6). Taken together, these results suggest that the 5’-UTR of A3G mRNA folds into independent domains and that mutations did not significantly impact the RNA secondary structure.

### Initiation at the uORF is required in Vif-mediated A3G translation inhibition

To further characterize the molecular mechanism of Vif-mediated A3G translation inhibition, we sought to determine if the uORF plays a role in this translational repression. We therefore transfected various A3G mRNA mutants in HEK 293T cells in presence or absence of Vif and with or without the proteasome inhibitor ALLN in order to discriminate translational inhibition from proteasomal degradation (42). First, when we co-transfected wt A3G construct with Vif, we observed, as expected, a strong decrease in A3G expression due to a cumulative effect of proteasomal degradation and translation inhibition by Vif (Figure 4, red bar, see also Figure S1 for western blots), whereas in presence of ALLN, we observed a typical 30-40% reduction in A3G synthesis due to A3G translational repression by Vif (Figure 4, blue bar), as previously observed (42). In a second step, we similarly analyzed the effect of Vif (+/−ALLN) on different A3G mRNA constructs. The results showed that A3G ΔuORF mutant presents a strong inhibition of A3G protein expression in presence of Vif (between 60-70%) (Figure 4, mutant ΔuORF), but less pronounced than with the wt A3G construct. Furthermore, in presence of ALLN, we did not observe any significant decrease in A3G expression when Vif was present (Figure 4, blue bars), indicating that the uORF is required for the Vif-mediated translational inhibition. To validate this finding, we transfected the A3G suAUG construct containing a single substitution at the upstream initiation codon. As expected, translational inhibition was not observed with this mutant (Figure 4, blue bar), and inhibition in the absence of ALLN was also reduced compared to the wt construct (Figure 4, red bar).

**Figure 4:**
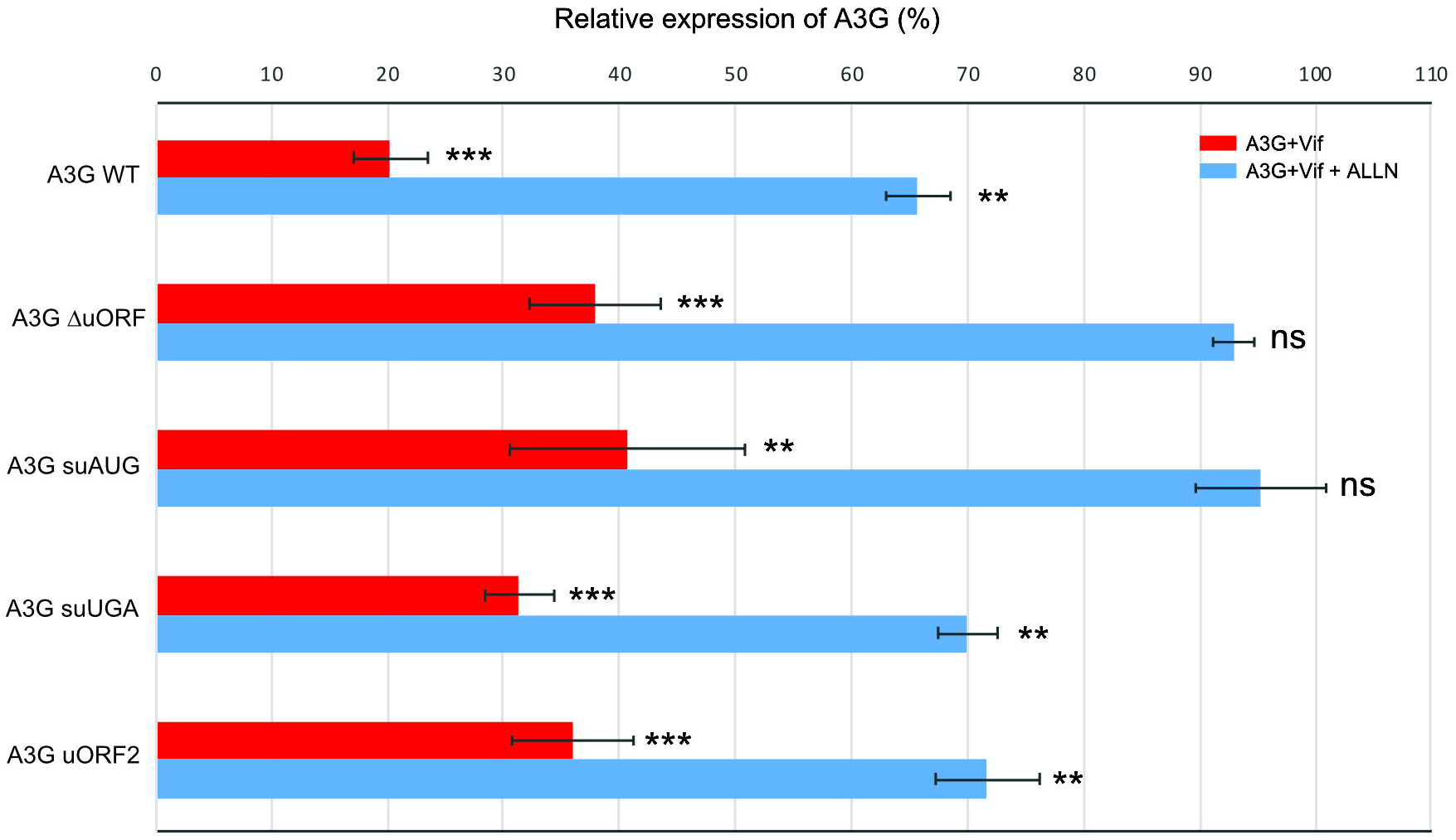
Effect of the uORF on Vif-mediated A3G translation inhibition. HEK 293T cells were transfected with plasmids expressing wt and mutated A3G constructs in the presence or absence of Vif and in the presence or absence of a proteasome inhibitor (ALLN). Proteins were separated by SDS/PAGE and analyzed by immunoblotting (see supporting Figure 2). Bands were quantified using Image J, normalized to actin, and relative expression of A3G proteins is represented. Histograms represent the effects of Vif on A3G: degradation + translation (red bars) and translation only (blue bars). Each condition is compared to the corresponding one without Vif and set to 100% (as above). Data represent the mean ± S.E.M. for at least three independent experiments. P-values are indicated as follows: *<0,05; **<0,01; ns: not significant.

Finally, we asked whether the inhibition of re-initiation (A3G suUGA) and the peptide identity (A3G uORF2) were important for the translational inhibition of A3G by Vif. These two constructs were transfected into HEK 293T cells as described above and analyzed by western blot (Figure S1). As expected, A3G proteins expressed from these constructs are degraded through the proteasome in presence of Vif (Figure 4, red bars). Interestingly, both mutants presented a wt A3G expression level in the presence of Vif and ALLN (Figure 4, blue bars), suggesting that (i) the uORF peptide has no role in the Vif-induced translational inhibition (A3G uORF2), and (ii) Vif only affects the leaky scanning (re-initiation is inhibited in the A3G suUGA mutant). Of note, none of the mutations significantly impacted RNA expression levels (Figure S2). Taken together, these results indicate that translation initiation at the uORF is essential for the Vif-mediated A3G translational inhibition.

### Relocation of A3G mRNA to stress granules is dependent on the uORF and Vif

Mechanisms of translational control dictate which mRNA transcripts gain access to ribosomes, and this process is highly regulated by the interplay of RNA binding proteins (RBPs) and RNA granules, such as processing bodies (P-bodies) or stress granules (SG). SGs are transient foci enriched in translation initiation factors and 40S ribosomal subunits, whereas P-bodies are enriched in RNA-decay machinery. Hence, SGs and P-bodies can be considered as extensions of the messenger ribonucleoprotein (mRNP) translational control cycle, i.e. as compartments where translationally silenced mRNPs are stored (66, 67). Thus, we asked whether Vif can relocate A3G mRNA into these storage compartments and participate to the downregulation of A3G translation, and if so if the uORF plays a role in this process. To test this hypothesis, we analyzed the wt A3G mRNA and two mutant constructs: A3G Δ5’-UTR and A3G suAUG. These two mutants (deletion of the 5’-UTR and single substitution of the uAUG) have been chosen because their translation is not down-regulated by Vif ((42) and this study). We began by a careful examination of the intracellular localization of A3G mRNAs and proteins by FISH and immunofluorescence (IF) analysis, respectively, after transfection of HEK 293T cells Figure S3). As expected, wt A3G mRNA and protein were detected in the cytoplasm (Figure S3, lane 4). Similarly, mRNAs and proteins expressed from mutants A3G Δ5’-UTR and suAUG were also present in the cytoplasm (Figure S3, lanes 5 & 6), suggesting that mutations do not impact mRNA localization and protein expression.

Next, we analyzed the co-localization of A3G mRNAs and proteins with SG and P-body markers in presence or absence of Vif. We co-transfected HEK 293T cells with A3G constructs in presence or absence of Vif, and then briefly exposed them to arsenite sodium (ARS) or high temperature (44°C) to induce a stress condition. We then performed FISH analysis (A3G mRNAs) and immunofluorescence staining using antibodies directed against Vif and A3G. Stress granule (PABP1 and TIA-1) and P-body (AGO2 and DCP1) marker proteins were expressed as GFP fusion proteins, allowing their direct fluorescence detection. First, concerning wt A3G mRNA, we observed as expected that PABP1 was localized into SGs regardless of stress conditions (Figure 5A). Interestingly, under stress conditions, while we observed a clear co-localization of A3G and PABP1 proteins into punctate granules in presence or absence of Vif (Figure 5A, column 7), co-localization of A3G mRNA and PABP1 was mainly observed in presence of Vif (Figure 5A, column 6, yellow dots; Figure S4). We obtained between 50-70% and 45-55% co-localization for PABP1/A3G mRNA (Figure 5D, blue bars) and TIA-1/A3G mRNA (Figure 5E, blue bars), respectively, in presence of Vif. Meanwhile, we also performed FISH and IF analysis for A3G Δ5’-UTR (Figure 5B) and suAUG (Figure 5C) mRNA mutants. Interestingly, whereas a co-localization of A3G and PABP1 proteins was still detected for these two constructs (Figure 5B & C, column 7; Figure S3), we observed a significant decrease (around 30%) of A3G Δ5’-UTR (Figure 5D red bars) and A3G suAUG (Figure 5E, green bars) mRNAs into SGs in the presence of Vif, suggesting that relocation of A3G mRNA into SGs depends not only on Vif but also on the uORF functionality. Of note, stress conditions (ARS or 44°C) did not relocate the different A3G mRNAs in the absence of Vif (Figure 5D & 5E, left part). Finally, we performed similar experiments with P-bodies markers (AGO2 and DCP1) (Figure 6). Under physiological conditions, while A3G and AGO2 proteins co-localized (Figure 6A, column 7), A3G mRNAs (wt or mutants) were rarely observed co-localizing with P-bodies markers (less than 20% co-localization), regardless of the presence of Vif (Figure 6B).

**Figure 5:**
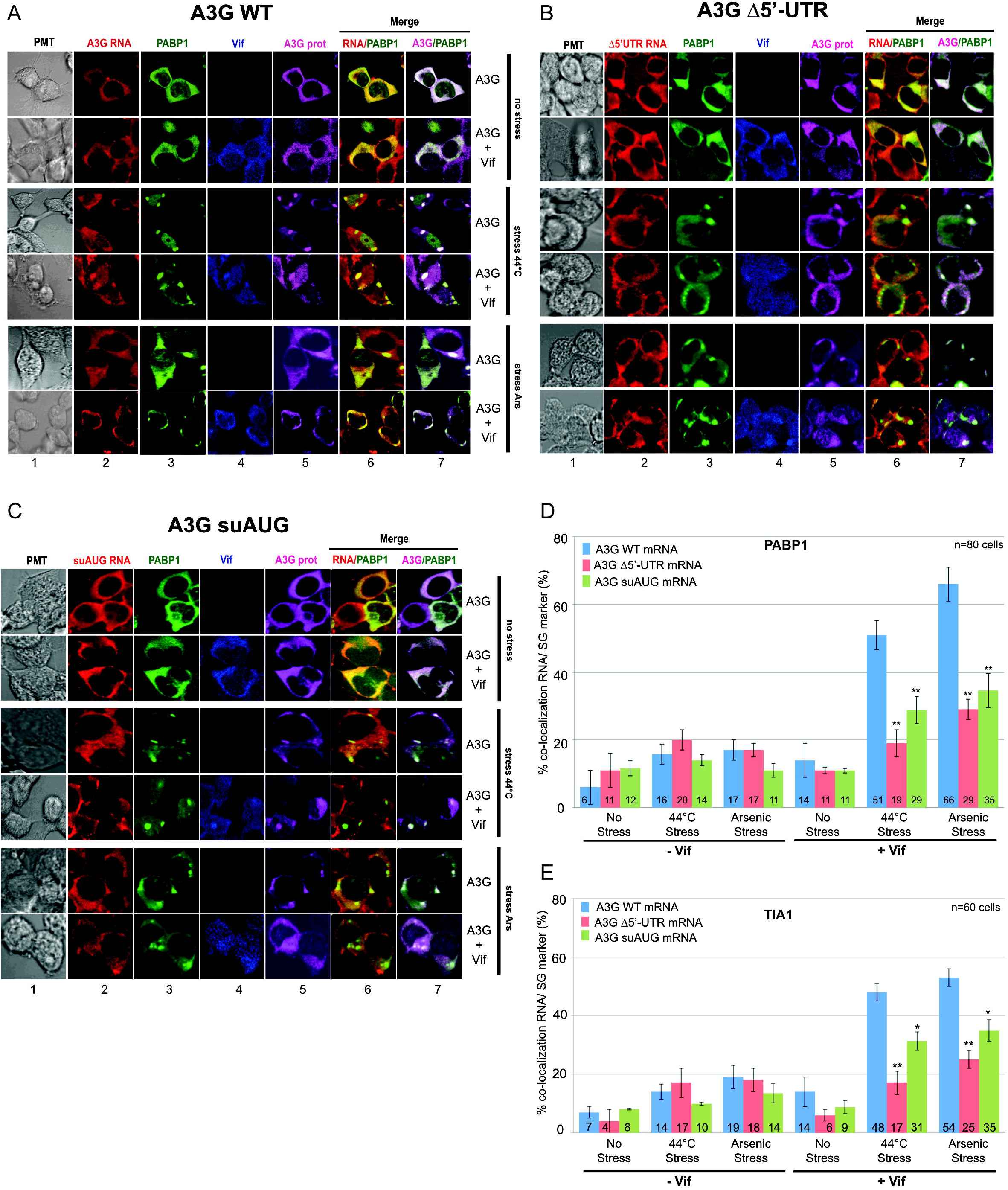
Importance of the uORF for the relocation of A3G mRNA into stress granules by Vif. HEK 293T cells were transfected with plasmids expressing wt A3G (**panel A**) as well as Δ5’-UTR (**panel B**) and suAUG (**panel C**) mRNAs, in absence or presence of Vif, and with a vector expressing GFP-PABP1 (SG marker). Cells were cultured in various conditions: (i) untreated (no stress) or stressed by (ii) incubation at 44°C or (iii) with arsenite sodium (Ars). Cells were fixed and probed with anti-DIG (A3G mRNAs), anti-A3G (A3G protein), and anti-Vif antibodies. PABP1 and SGs were visualized by direct fluorescence of the GFP-PABP1 fusion protein. Cells were stained with Dapi to visualize nuclei and the images were merged digitally. **D**) Histograms represent the percentage of co-localization of A3G mRNAs with PABP1 or with TIA (**E**). Standard deviations are representative for at least three independent experiments. P-values are indicated as follows: **<0,01.

**Figure 6:**
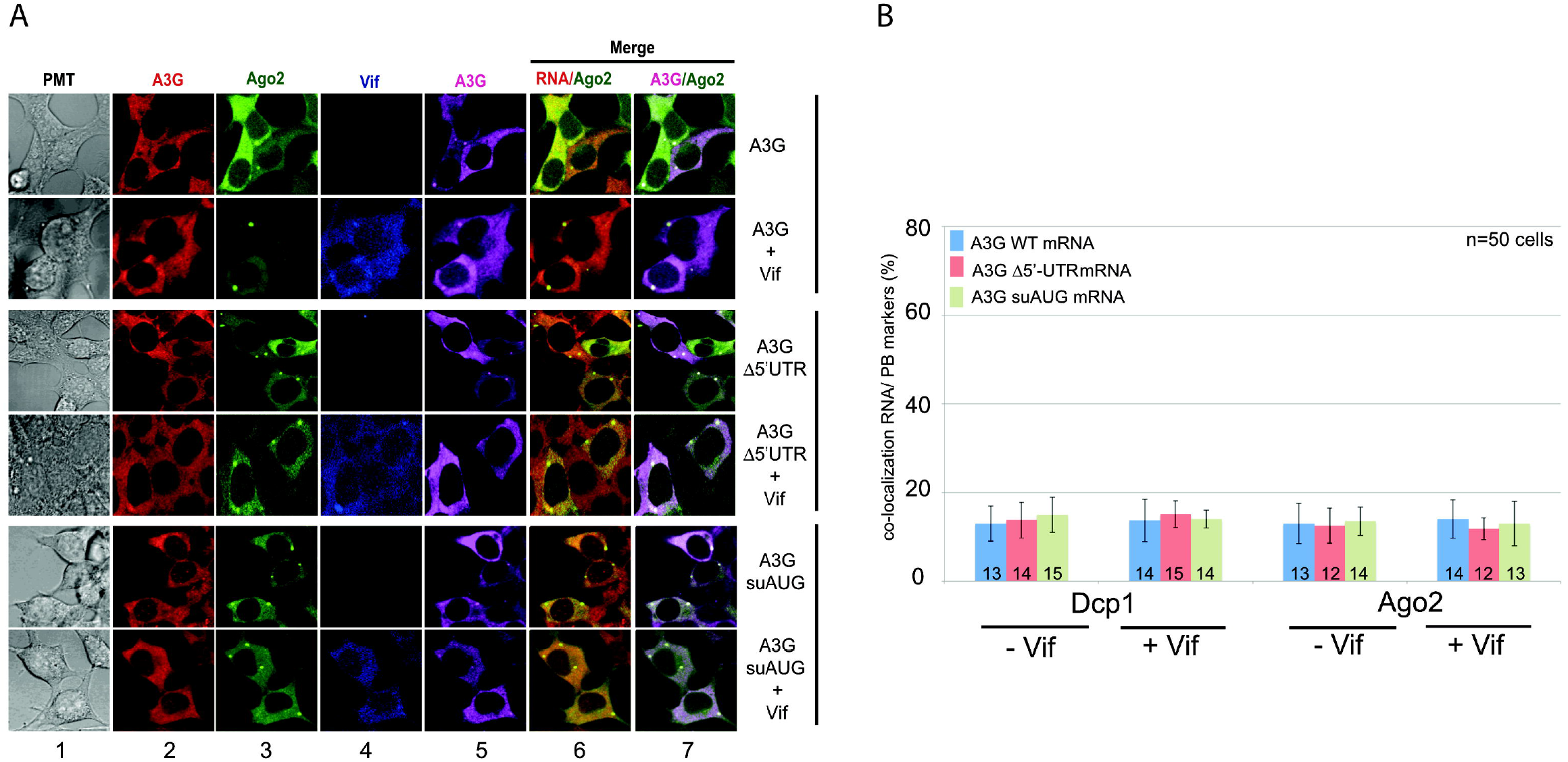
The uORF does not contribute to A3G mRNA relocation to P-bodies by Vif. **A)** HEK 293T cells were transfected with plasmids expressing wt A3G as well as Δ5’-UTR and suAUG mRNAs, in absence or presence of Vif, and with a vector expressing GFP-AGO2 (P-body marker). Cells were fixed and probed with anti-DIG (A3G mRNAs), anti-A3G (A3G protein), anti-Vif antibodies. AGO2 was visualized by direct fluorescence of the GFP-AGO2 fusion protein. Cells were stained with Dapi to visualize nuclei and the images were merged digitally. B) Histograms represent the percentage of co-localization of A3G mRNAs with AGO2 or DCP1 (another P-body marker). Standard deviations are representative for at least three independent experiments.

## DISCUSSION

The HIV Vif protein acts as a key antagonist of A3G and as such it plays a major role in promoting efficient viral infection. Vif counteracts A3G according to two distinct mechanisms that concur in reducing its intracellular levels: direct protein degradation and protein translation inhibition. While mechanisms leading to A3G degradation through the recruitment of an E3 ubiquitin ligase complex by Vif are well understood, little is known concerning its translational regulation by Vif. Recently, we showed that the 5’-UTR of A3G mRNA, and specifically the SL2-SL3 domain, was required for the Vif-induced translation inhibition of A3G (42). In the present study, we report that a highly conserved uORF embedded within these structures plays a crucial role in modulating A3G translation both under physiological conditions and during Vif-mediated translation repression.

Here, we showed that translation of A3G was significantly increased when the uORF was disrupted (deletion of the uORF or substitution of the uAUG) (Figure 3), indicating that the uORF is indeed a repressive element, as previously observed for several other genes like the tyrosine kinases HCK (Hematopoietic Cell Kinase), LCK (Lymphocyte specific protein tyrosine Kinase), ZAP70 (Zeta chain of T cell receptor Associated Protein 70 kDa), YES1 (yes proto-oncogene 1, Src family tyrosine kinase) or the oncogenes MDM2 (Murine Double Minute 2) and CDK4 (Cyclin Dependent Kinase 4) (68). Generally, the presence of uORFs modulates ribosome access to the downstream ORF and can reduce its translational efficiency by 30–48% (54). While current data do not suggest that A3 are oncogenes, recent literature clearly showed that A3A and A3B are active on cancer genomes and may impact tumor progression (69). Previous studies found that A3G is overexpressed in patients with diffuse large B-cells leukemia (70) and in pancreatic cancers (71). While cancer development is multifactorial, it is tempting to speculate that cells evolved to limit their expression and that the uORF in the 5’-UTR of A3G mRNA may participate to this repression/regulation. Endogenous retroelements, such as ERV, LINE-1, or Alu elements (∼42 % of the human genome) (72, 73) are also finely regulated to avoid genetic diseases and cancers (74, 75). Interestingly, A3 proteins have evolved to protect hosts from the genomic instability caused by retroelements (76) and have been shown to counteract LINE-1 and Alu retrotransposition (77–83). Translational control of mRNA is an important feature of innate immunity but even if the mechanisms remain ill-defined, fine-tuning of their expression must be advantageous for the cell integrity.

uORFs can regulate translation by multiple mechanisms (84, 85). Using mutated A3G mRNA constructs to analyze the impact of the uORF on these various mechanisms, we showed that (i) the distance between the 5’ cap and the uORF does not impact A3G translation (42); (ii) the context around the uAUG is favorable to initiate translation, indicating that the uAUG is efficiently recognized by the scanning ribosomes but can also be leaky-scanned by the ribosome which will then initiate at the main AUG to express A3G; (iii) A3G may also be translated through a re-initiation mechanism as lengthening the uORF or shortening the intercistronic distance enhances translation repression (86–88). In this later case, and as previously observed, the distance between the uORF stop codon and the initiation codon of the main ORF would be too short for the scanning 40S to reacquire a new eIF2•GTP•Met-tRNA^iMet^ complex and initiate translation at the main AUG (89–91). Surprisingly, we did not observe any increase in A3G expression when we reduced the size of the potentially encoded uORF peptide (Figure 3). Indeed, these mutants present a longer intercistronic region (Figure 3A) and should have benefit from re-initiation. While this mechanism is not fully understood yet, it is possible that the intrinsic sequence of the uORF or its secondary/tertiary structure is of importance for this repression which might be caused by stalling of the elongating ribosomes. While we did not assess the densities of ribosome profiling reads over the 5’UTR of A3G mRNA, we retrieved data from the “Genome Wide Information on Protein Synthesis” website (https://gwips.ucc.ie/) and could observe that the uORF is occupied by ribosomes (Figure S7A). Interestingly, ribosome occupancy is higher in the second half of the uORF. This suggests the ribosomes stall during elongation and create a roadblock, hindering the scanning 43S preinitiation complexes that pass through the uORF start codon, which would explain the reduced translation of A3G (a similar profile could be observed for A3F 5’-UTR, Figure S7B). Besides, we showed that the identity of the putative 23 amino acids peptide expressed from the uORF (mutant A3G uORF2) has no function in the translational regulation of A3G, reinforcing the importance of the uORF (sequence/structure) over the peptide sequence.

Furthermore, we showed that the highly structured 5’-UTR of A3G mRNA ((40) and Figures S5, S6) did not drive significant translation through a potential IRES, as the insertion of a stable stem-loop at the 5’-end of the mRNA designed to prevent 43S subunit loading, almost completely inhibited A3G translation (Figure 3). Moreover, a genome-wide search yielding a large number of mammalian cellular IRESs did not identify the 5’-UTR of A3G as potential IRES (92). Altogether, these results suggest that A3G is translated through a dual leaky-scanning and re-initiation mechanism. Further studies will be needed to evaluate: (i) the re-initiation efficiency; accurate determination will be difficult as differentiating sole re-initiation from a leaky-scanning/re-initiation mechanism is hard to achieve; (ii) how the ribosome is stalled, through a specific RNA structure or the termination context (93).

Beyond effects of the uORF on A3G translation, studies in mammalian cells and yeast showed that uORF-containing mRNAs are susceptible to be targeted by NMD, which is attributed to the termination events occurring at uORF stop codons (94, 95). Moreover, the presence of uORF in the 5’ leader of mRNAs was associated with reduced transcript level and reduced translation, which together resulted in a decrease in ORF translation efficiency (averaging 30-48% reduction) (54). Indeed, under conditions where the proteasomal pathway was inhibited, we showed that Vif was not able to reduce the translation of A3G expressed from mRNA constructs devoid of a functional uORF (ΔuORF and suAUG), suggesting that Vif induces A3G translational repression during the initiation steps at the uORF, or that it increases stalling of elongating ribosomes. To the best of our knowledge, the uORF of A3G mRNA would represent the second example (and the first one in vertebrates) for a uORF being co-regulated by a protein factor. The first example being SXL (sex lethal protein), which mediates translation inhibition through association to uORFs of the Drosophila msl2 5’-UTR mRNA (96–98). Indeed, in females, msl-2 expression needs to be repressed for viability (dosage compensation), and this repression is achieved by the binding of SXL to uridine stretches in the 5’ and 3’UTRs (99, 100). Both mechanisms synergize to achieve full msl-2 translational repression (96). Similarly, multiple mechanisms also exist to inhibit A3G expression through its proteasomal degradation and translational inhibition by Vif. Whereas additional experiments will be needed to clearly decipher this mechanism, one can imagine that Vif interacts with components of the eukaryotic translation initiation machinery to reduce the translational rate of A3G or may recruit cellular factors involved in the negative regulation of the translation (Figure 7).

**Figure 7:**
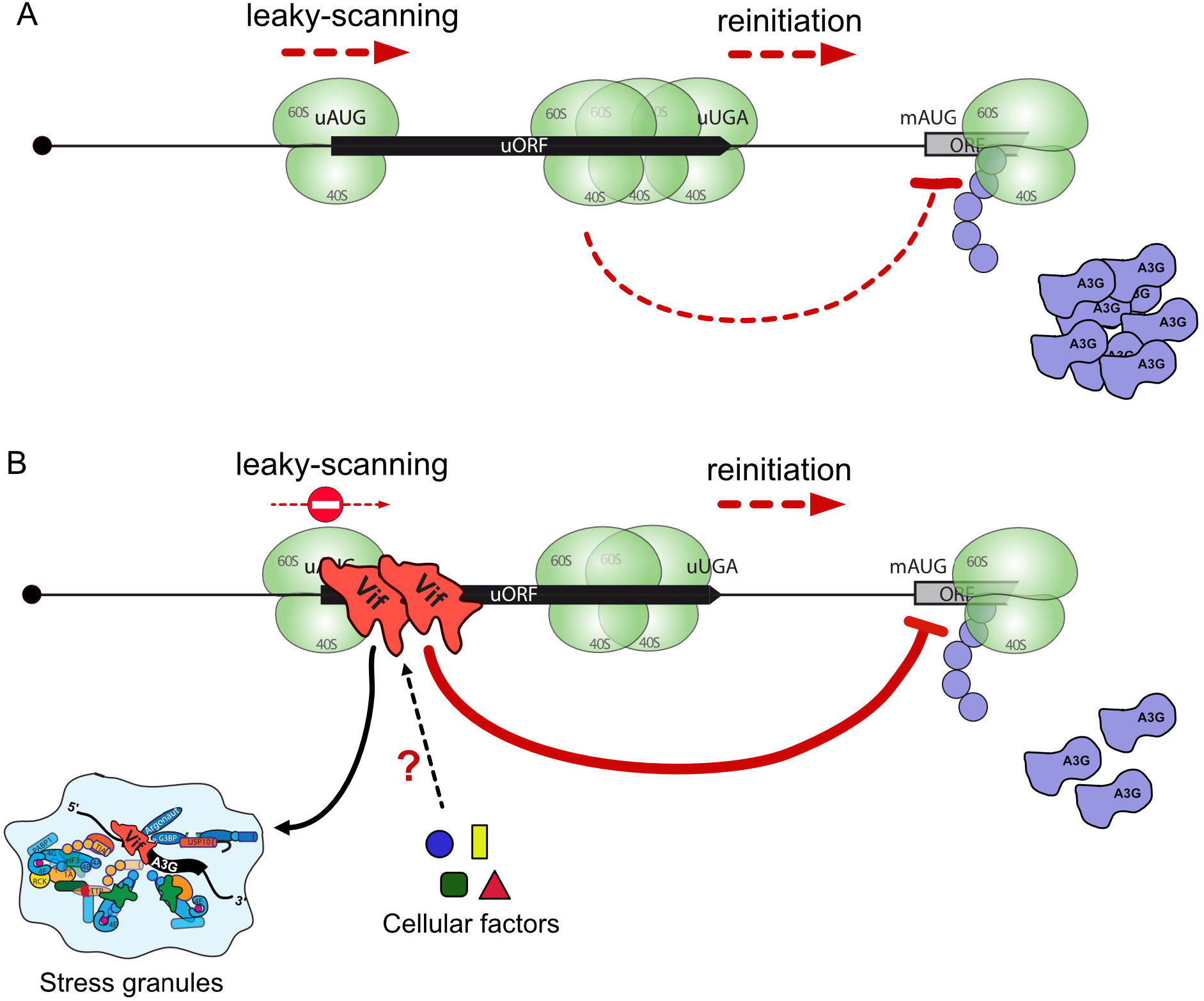
Simplified model of A3G mRNA uORF-mediated translational control. **A)** In absence of Vif, translation of A3G mRNA is controlled by the uORF embedded within its 5’UTR, leading to basal A3G protein translation through leaky-scanning and reinitiation. Ribosome stalling at uORF termination codon is probably participating in reducing A3G yield. **B)** In presence of Vif, the uORF strengthens the translational inhibition leading to reduced yield of A3G protein, mainly by inhibiting the leaky-scanning mechanism. Vif may interact with components of the eukaryotic translation initiation machinery to reduce the translational rate of A3G or recruit cellular factors involved in the negative regulation of the translation. Moreover, relocation of A3G mRNA into storage compartments in presence of Vif participates to the global reduction of A3G level.

Finally, we observed a strong correlation between the presence and functionality of the uORF on A3G mRNA and its co-localization, under stress conditions, into SGs in a Vif-dependent manner. Indeed, when the uORF was not functional (inhibition of uORF initiation or ΔuORF), the presence of mutated A3G mRNAs was significantly reduced into SGs when Vif was co-expressed (Figure 5). However, A3G protein expressed from these constructs can be found in P-bodies and SGs, as previously observed (57, 101–103). The presence of wt A3G mRNA in SGs in presence of Vif is probably not so surprising, since SGs play an important role in the regulation of gene expression at the translational level in response to a variety of external stimuli (104). Therefore, these results validate the notion that the uORF acts as a negative regulator of A3G expression by directing the relocation of A3G mRNA into storage compartments in the presence of Vif (Figure 7).

To summarize, we have identified a short uORF within the 5’-UTR of A3G mRNA that is conserved in the human population, regulates A3G translation and is required by HIV-1 Vif to repress its translation (Figure 7). While the relocation in SGs of A3G mRNA by Vif through the uORF could explain in part the downregulation of A3G expression, additional work will be required to firmly establish this mechanism. Deciphering the mechanisms of the Vif-mediated translational inhibition of A3G mRNAs will be important to find new molecular inhibitors able to counteract Vif activity and reduce viral infectivity.

## ACKNOWLEDGEMENT

We thank Drs Vincent Caval (Institut Pasteur Paris, France) and Sébastien Pfeffer (IBMC-CNRS, Strasbourg) for kindly providing the APOBEC3A and the different P-body and SG markers, respectively. The following reagents were obtained through the AIDS Research and Reference Reagent Program, Division of AIDS, NIAID, NIH: anti-A3G polyclonal antibody (#9968) from Dr Warner C. Greene, HIV-1 Vif monoclonal antibody (#319) from Dr Michael H. Malim, rabbit anti-human A3F polyclonal antibody (#11226) from Immunodiagnostics, and anti-human A3H monoclonal antibody (P3A3-A10, #12155) from Drs Michael Emerman and Reuben Harris.

## AUTHOR CONTRIBUTIONS

JCP designed and supervised the project. CL, TS, SG, BS and MW constructed and characterized all A3G mutants by cell transfection and immunoblotting. OG, CV and RCR performed the RACE, qRT-PCR, and SHAPE, respectively. JB performed FISH and IF experiments. LE performed the genetic analyses of A3G/A3F genes. JCP, RM, AC, and LE analyzed the data. JCP wrote the manuscript with contributions from the other authors. All authors have read and agreed to the published version of the manuscript.

