## Supplementary figure legends for "A conserved uORF regulates APOBEC3G translation and is targeted by HIV-1 Vif protein to repress the antiviral factor"

**Legends to supplementary figures and tables**

**Figure S1: Effect of Vif on the translation of the different A3G mRNA constructs**. HEK 293T cells were transfected with plasmids expressing wt and mutated A3G constructs in the presence or absence of Vif and in the presence or absence of a proteasome inhibitor (ALLN). Proteins were separated by SDS/PAGE and analyzed by immunoblotting. Bands were quantified using Image J and relative expression of A3G proteins are represented in histograms (see Figure 4).

**Figure S2: A3G mRNA expression level in HEK 293T transfected cells.** Total RNA was extracted from transfected HEK 293T cells and A3G RTqPCR were performed to study the expression of wt and mutated A3G constructs. Data represent the mean ± S.E.M. for at least three independent experiments.

**Figure S3: Analyses of A3G mRNAs and protein by FISH and immunofluorescence.** HEK 293T cells were transfected with plasmids expressing wt A3G as well as ∆5’-UTR and suAUG mRNAs. Cells were fixed and probed with anti-DIG (A3G mRNAs) and anti-A3G (A3G protein) antibodies. Cells were stained with Dapi to visualize nuclei and the images were merged digitally. Controls and A3G mRNAs (lines 1 to 6) are indicated on the right of the panels.

**Figure S4**: **Intensity plots (FISH)** obtained from the cytoplasmic signals of PAPB1 (green lines), A3G (pink lines), and A3G mRNA (red lines) in the absence or presence of Vif in physiological (**A**), or in stress conditions: 44°C (**B**) or sodium arsenite (**C**).

**Figure S5: Comparison of the SHAPE reactivity profiles of the 5’ region of wild-type and mutant A3G mRNAs**. The SHAPE reactivity profiles of nucleotides 1-400 for A3G mRNAs WT, suAUG, suUGA, WK, suUGA276 and suUGA289 are represented. The SHAPE reactivity values are drawn on the structure and color-coded as indicated in the inset. They were used as constraints for the RNAstructure software (version 6.0). Positions of uORF (red), translation initiation codons (uAUG for uORF and mAUG for A3G in blue), translation stop codon (uUGA for uORF in blue) and mutations (green) are indicated. Four independent domains can be deduced from these structures and are indicated on the wild-type structure.

**Figure S6: Comparison of the SHAPE reactivity profiles of the 5’ region of wild-type and mutant A3G mRNAs**. The SHAPE reactivity profiles of nucleotides 1-400 for A3G mRNAs WT, ∆uORF, ∆249-273 and ∆249-289 are represented. The SHAPE reactivity values are drawn on the structure and color-coded as indicated in the inset. They were used as constraints for the RNAstructure software (version 6.0). Positions of uORF (red), translation initiation codons (uAUG for uORF and mAUG for A3G in blue), translation stop codon (uUGA for uORF in blue) and mutations (green) are indicated. Four independent domains can be deduced from these structures and are indicated on the wild-type structure.

**Figure S7:** **Densities of ribosome profiling reads over the 5’UTR of A3G (A) and A3F (B) mRNAs**. Data were retrieved from the “Genome Wide Information on Protein Synthesis” website (<https://gwips.ucc.ie/>). Ribosomes profiles (red), mRNA-seq coverages (green), and position of the uORF (above the amino acids sequence) are indicated.

**Table S1**: **Variants identified in the human A3G and A3F uORF (+/- 10 nt) regions.** In bold, the common SNP (MAF>1%); In italics, variants that would impact uORF (*i.e*. at the uAUG position or inducing frameshift). Data from the NCBI dbSNP database (July 31^st^ 2019); Homo sapiens:GRCh38.p12 (GCF_000001405.38)Chr 22 (NC_000022.11).

**Table S2**: **5’-UTR sequences of APOBEC3 mRNAs identified by RACE-PCR.** APOBEC3 genes are indicated, as well as their 5’-UTR length. The translation initiation codon AUG of the main APOBEC3 ORF is indicated (blue) as well as the potential uORF (orange).

**Table S3: SHAPE data sets**. The different sheets correspond to SHAPE reactivities obtained for wt and mutated A3G mRNAs.
