## Supplementary figures and images for "A conserved uORF regulates APOBEC3G translation and is targeted by HIV-1 Vif protein to repress the antiviral factor"

### Figure S1

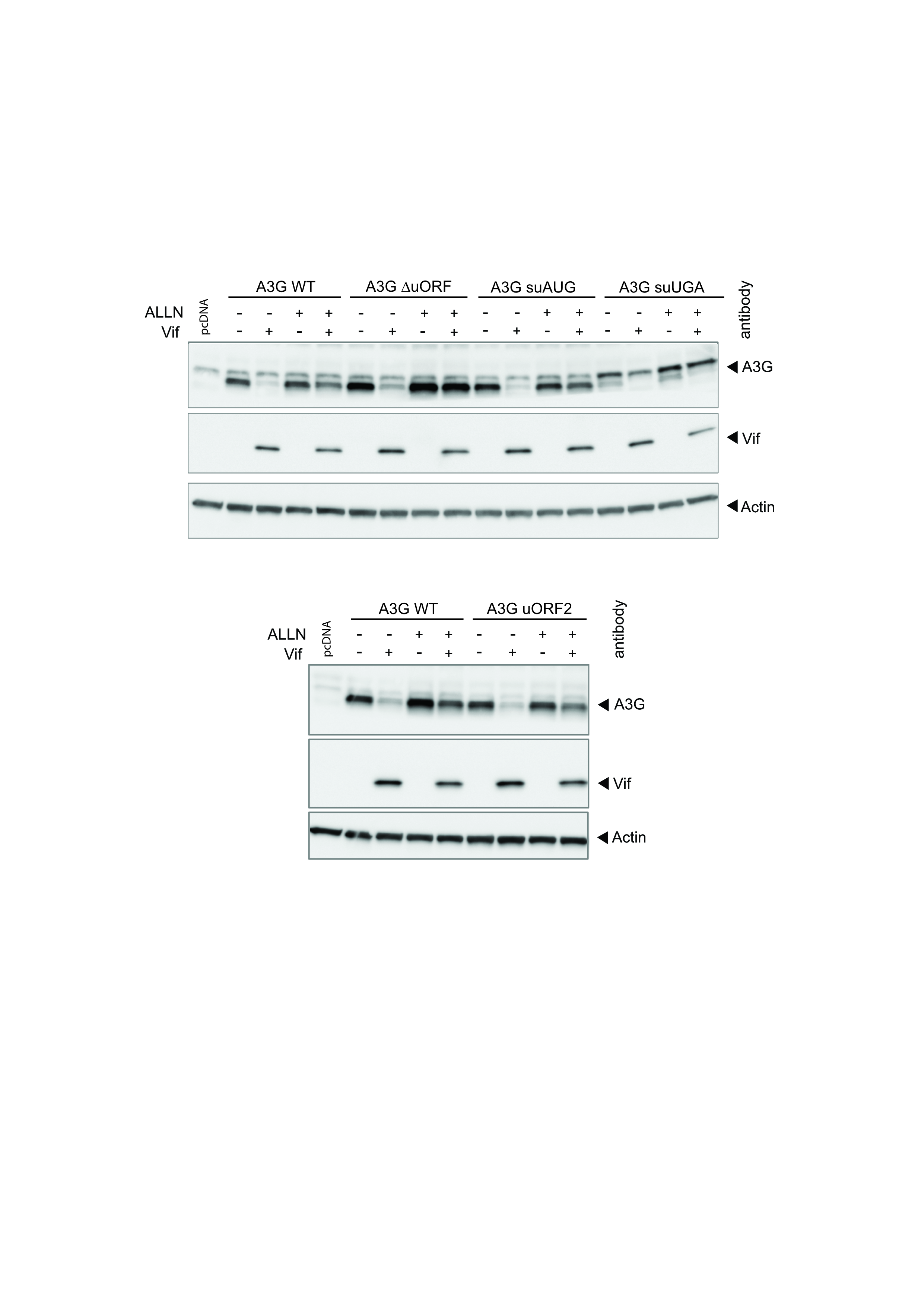

### Figure S2

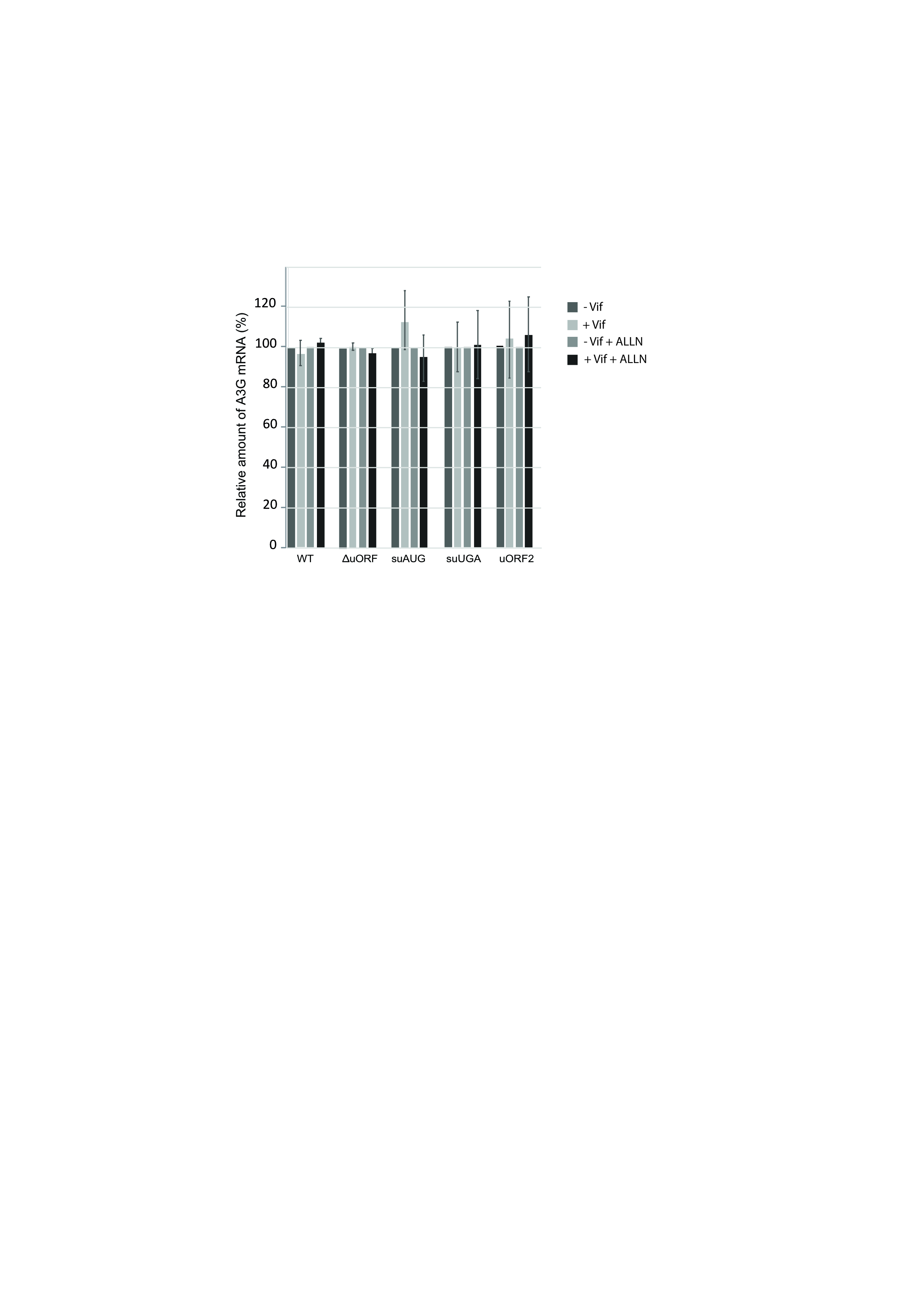

### Figure S3

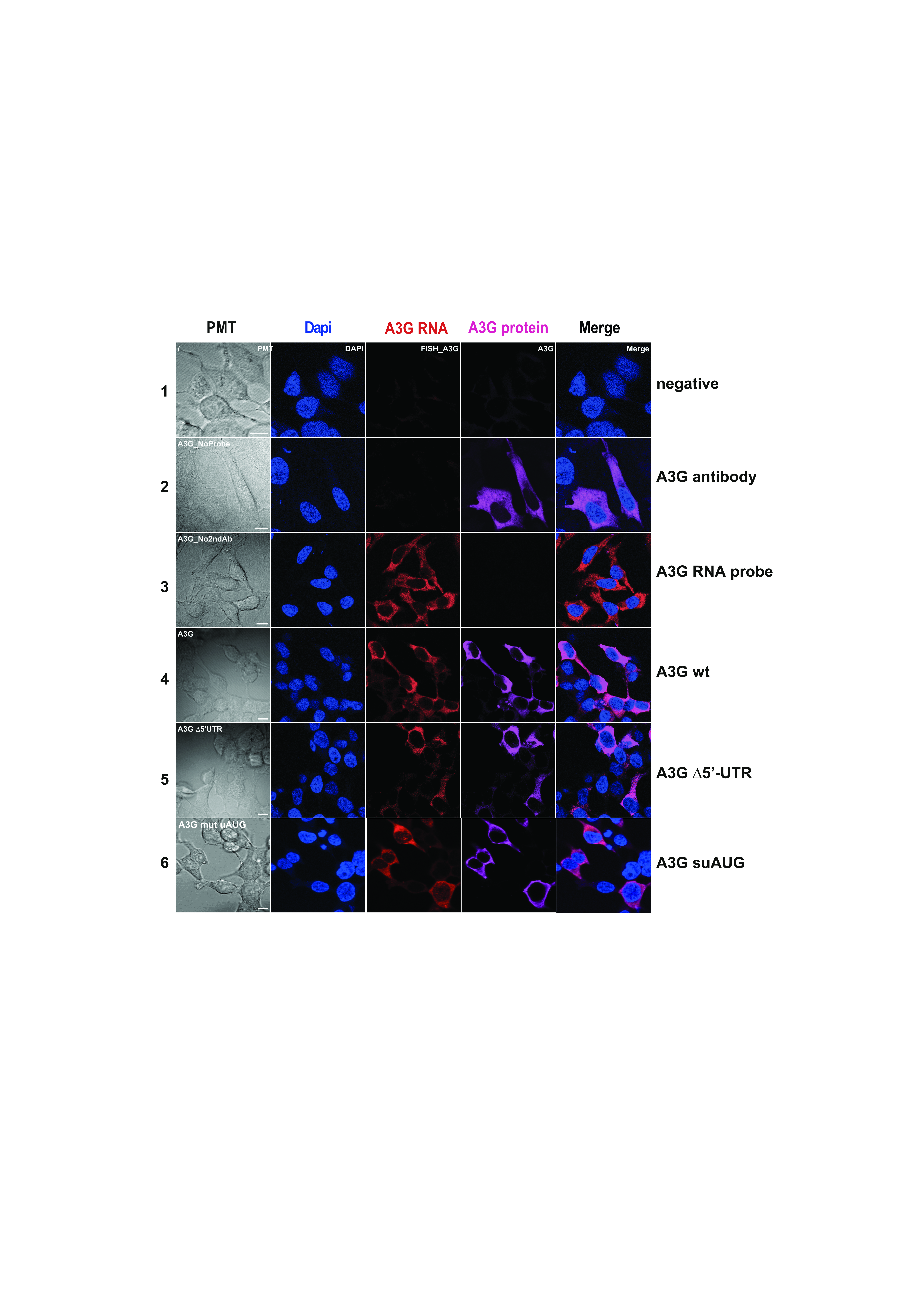

### Figure S4

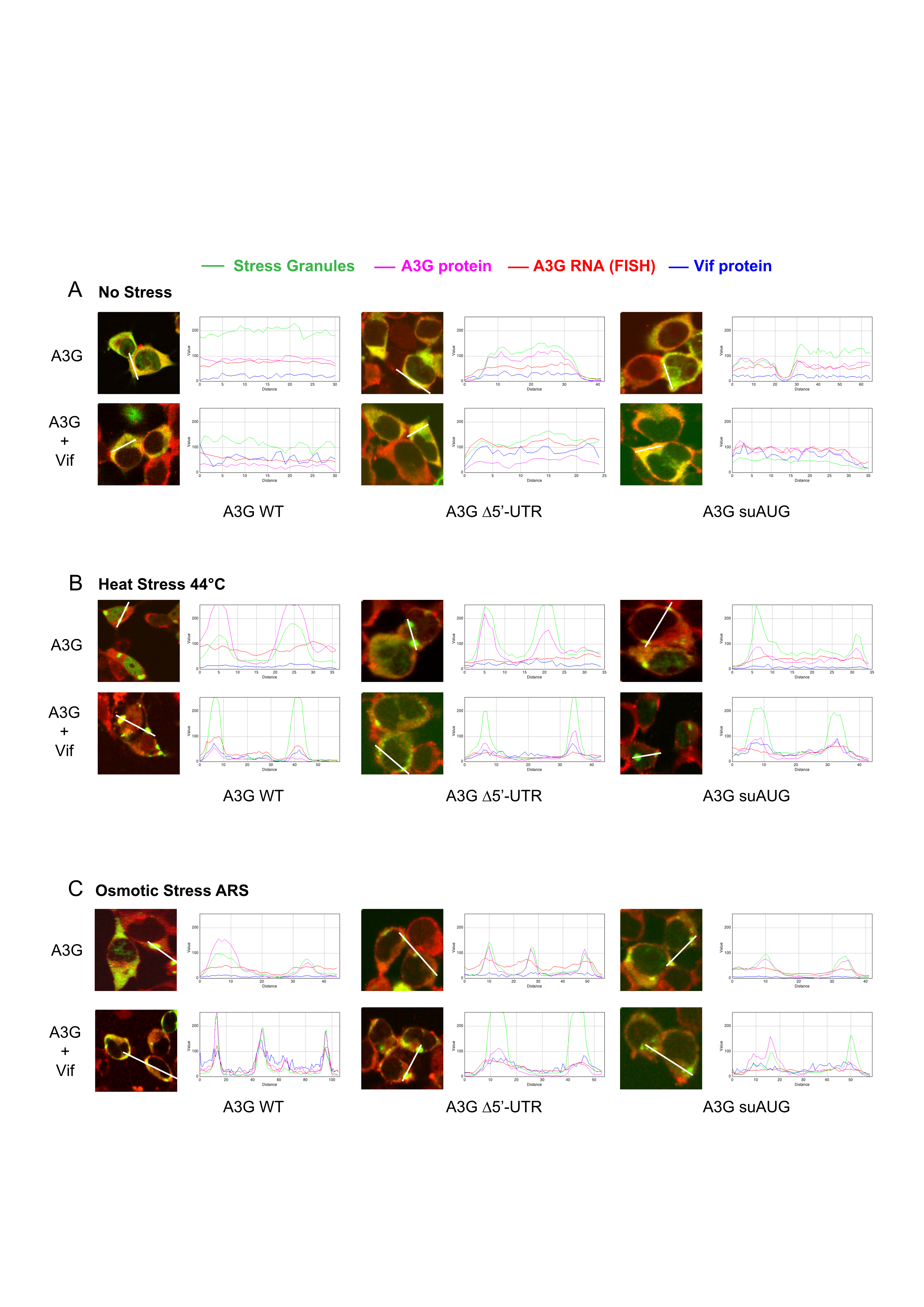

### Figure S5

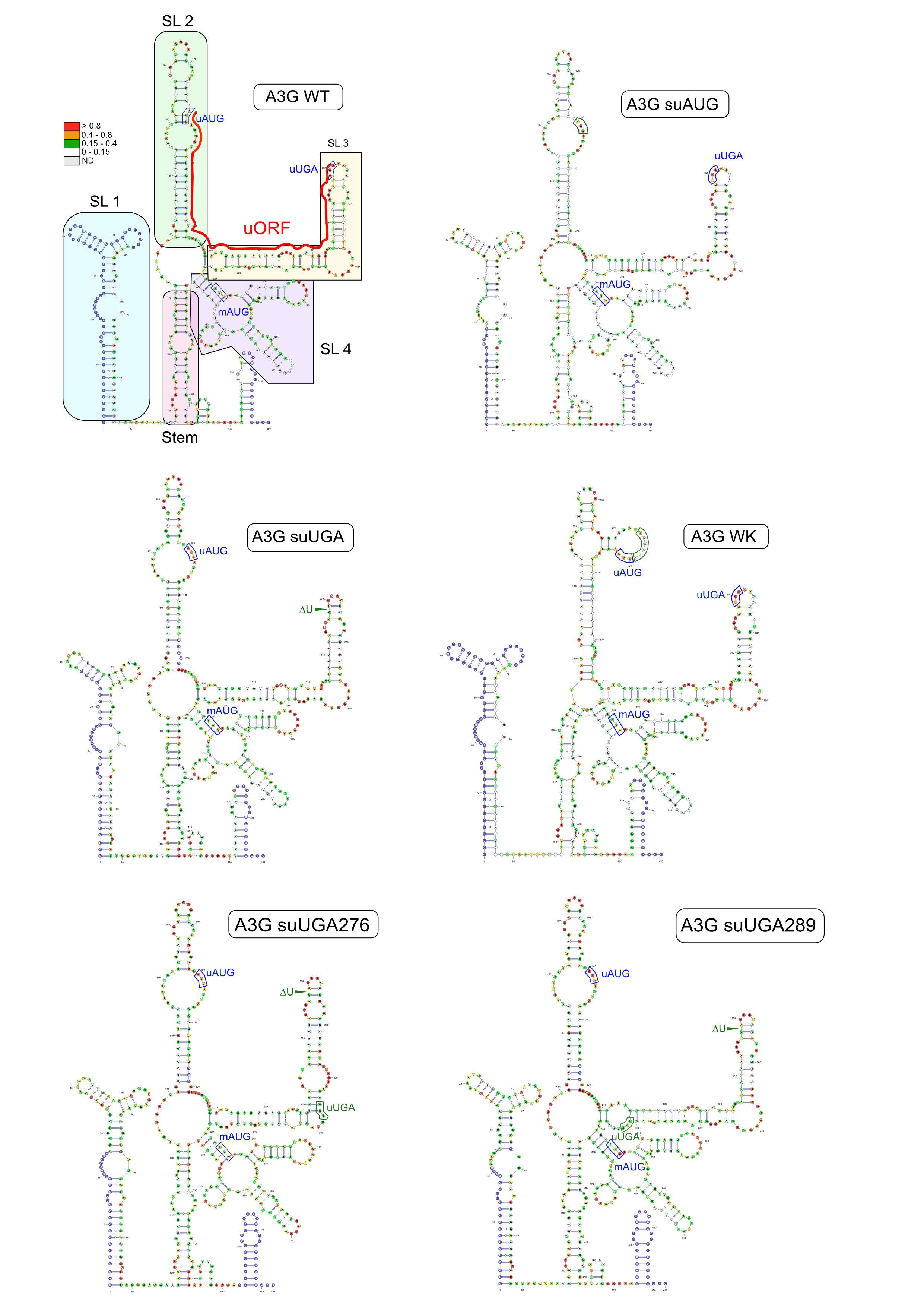

### Figure S6

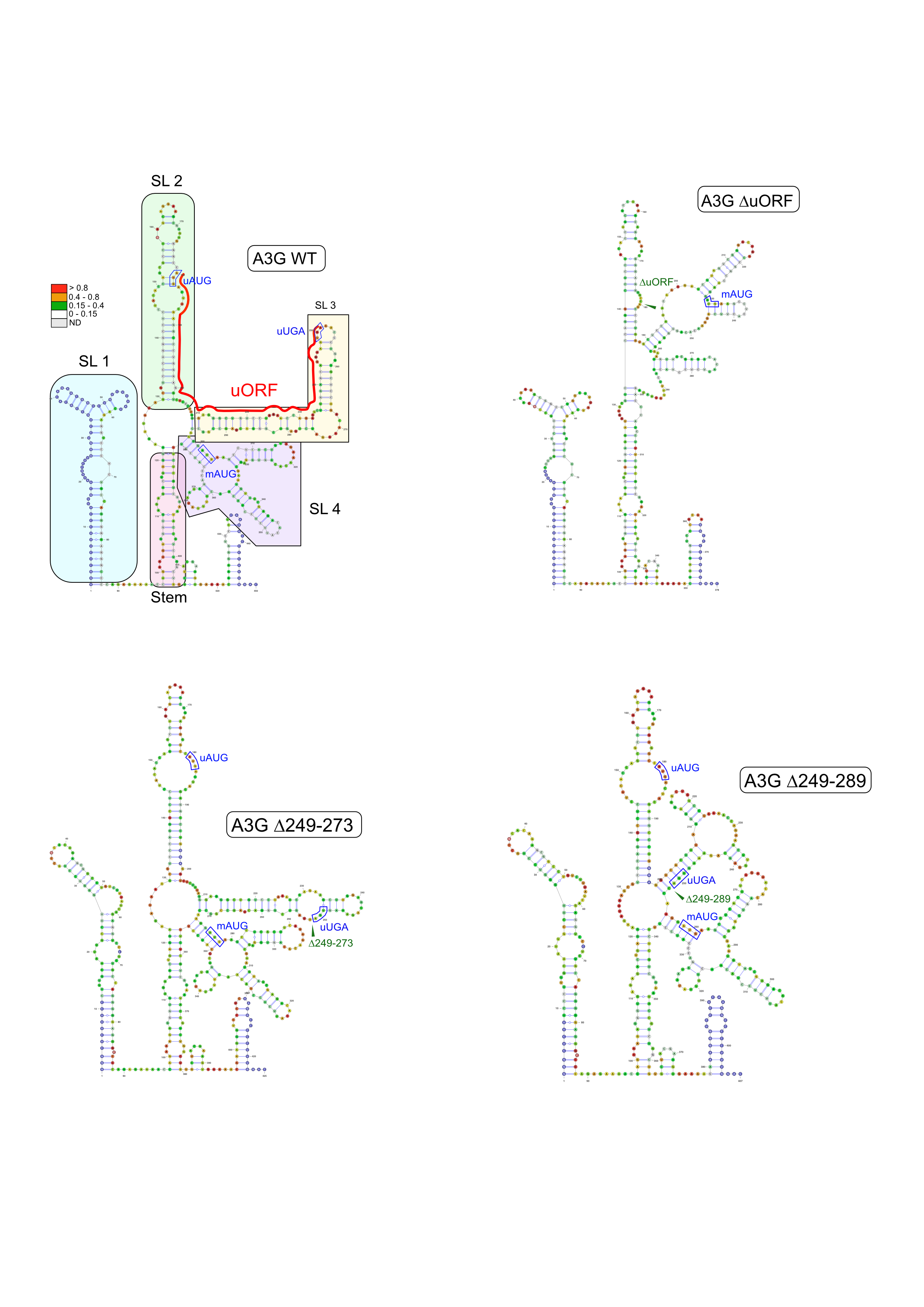

### Figure S7

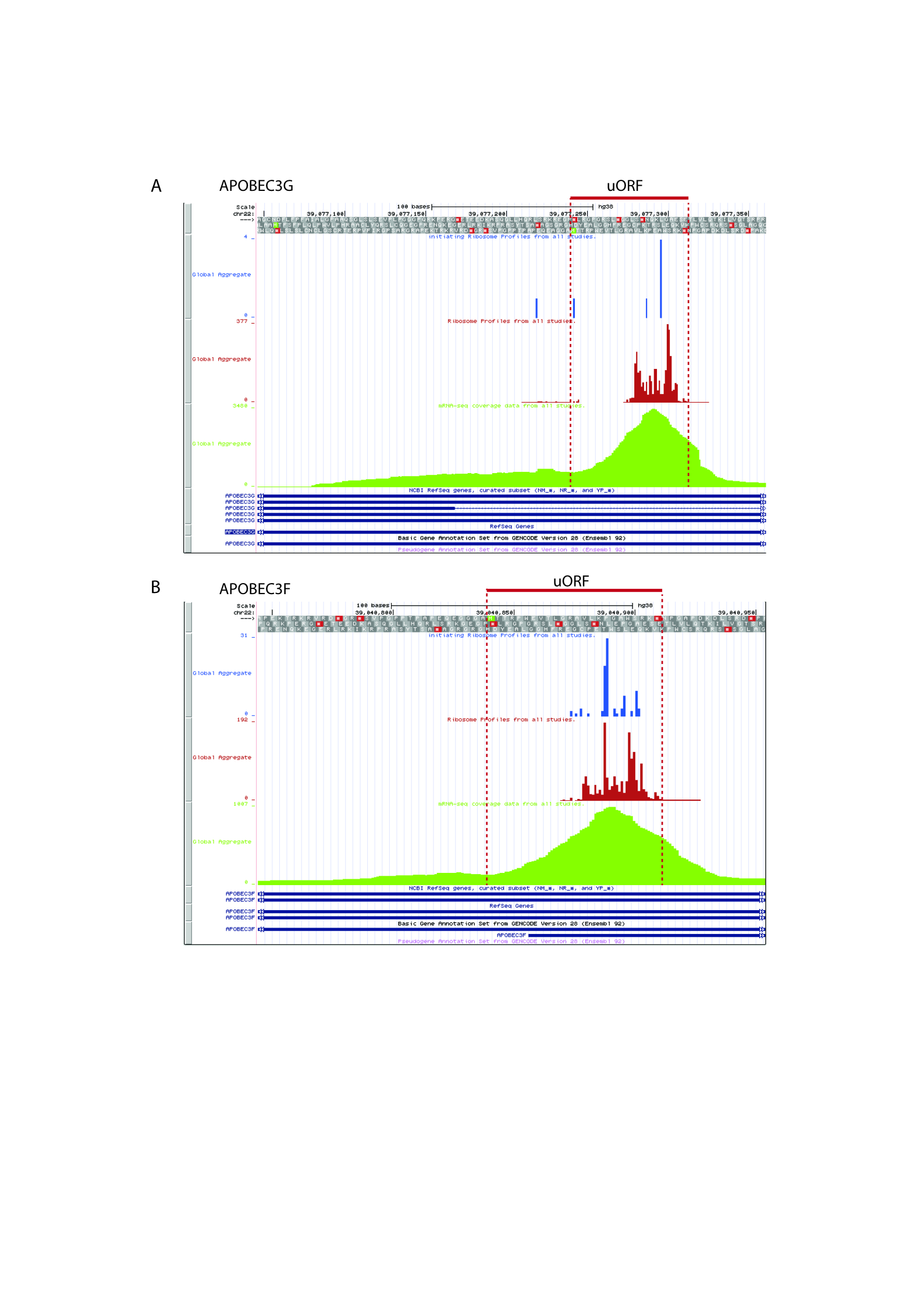
